# Decoding optimal ligand design for multicomponent condensates

**DOI:** 10.1101/2023.03.13.532222

**Authors:** Sarasi K. K. Galagedera, Thuy P. Dao, Suzanne E. Enos, Antara Chaudhuri, Jeremy D. Schmit, Carlos A. Castañeda

## Abstract

Biomolecular condensates form via multivalent interactions among key macromolecules and are regulated through ligand binding and/or post-translational modifications. One such modification is ubiquitination, the covalent addition of ubiquitin (Ub) or polyubiquitin chains to target macromolecules for various cellular processes. Specific interactions between polyubiquitin chains and partner proteins, including hHR23B, NEMO, and UBQLN2, regulate condensate assembly or disassembly. Here, we used a library of designed polyubiquitin hubs and UBQLN2 as model systems for determining the driving forces of ligand-mediated phase transitions. Perturbations to the UBQLN2-binding surface of Ub or deviations from the optimal spacing between Ub units reduce the ability of hubs to modulate UBQLN2 phase behavior. By developing an analytical model that accurately described the effects of different hubs on UBQLN2 phase diagrams, we determined that introduction of Ub to UBQLN2 condensates incurs a significant inclusion energetic penalty. This penalty antagonizes the ability of polyUb hubs to scaffold multiple UBQLN2 molecules and cooperatively amplify phase separation. Importantly, the extent to which polyubiquitin hubs can promote UBQLN2 phase separation are encoded in the spacings between Ub units as found for naturally-occurring chains of different linkages and designed chains of different architectures, thus illustrating how the ubiquitin code regulates functionality via the emergent properties of the condensate. We expect our findings to extend to other condensates necessitating the consideration of ligand properties, including concentration, valency, affinity, and spacing between binding sites in studies and designs of condensates.

**Highlights:** ● There is an optimal polyUb ligand architecture/design that promotes multicomponent phase separation, as polyUb hubs whose Ub units are too close together or too far apart are not effective drivers of phase separation for either UBQLN2 450-624 or full-length UBQLN2.

● Theoretical modeling reveals that Ub incurs a significant inclusion energetic penalty that is balanced by polyUb’s ability to act as a hub to amplify UBQLN2-UBQLN2 interactions that facilitate phase separation.

● Naturally-occurring M1-linked polyUb chains are optimized to maximize phase separation with UBQLN2.

● Different linkages used in the Ub code deliver biochemical information via Ub-Ub spacing, whereby different outcomes are regulated by the emergent properties of Ub-containing biomolecular condensates.

**Significance:** Biomolecular condensates are essential for cellular processes and are linked to human diseases when dysregulated. These condensates likely assemble via phase transitions of a few key driver macromolecules and are further modulated by the interactions with ligands. Previous work showed that ligands with one binding site inhibit driver phase transitions whereas ligand hubs comprising several identical binding sites to drivers promote phase transitions. Here, using a library of designed ligand hubs with decreasing or increasing spacings between binding sites and altered binding affinities with drivers, we employ theory and experiments to establish a set of rules that govern how ligand hubs affect driver phase transitions. Our findings reveal that effects of macromolecules can be manipulated through emergent properties of condensates.

## Introduction

Membraneless biomolecular condensates are an important mechanism for cells to quickly regulate molecular processes, such as stress response, gene expression, and protein degradation (1, 2). Biomolecular condensates comprise hundreds to thousands of unique macromolecules, but only a few of these are essential for the formation of condensates (3, 4). These essential components are classified as scaffolds whereas the nonessential components are known as clients/ligands. Scaffolds are multivalent macromolecules that can self-interact at specific sites called stickers to undergo liquid-liquid phase separation (LLPS) into coexisting dense and dilute liquid phases. In contrast, ligands, which do not phase separate on their own, bind to scaffolds and regulate condensate assembly/disassembly and material properties. Recent studies have uncovered several rules that dictate how different features of ligands can lead to modulation of scaffold phase behavior (5). Specifically, low- and high-valency ligands tend to inhibit and promote scaffold LLPS, respectively (6–13). Moreover, computational studies showed that weakly binding ligands do not affect scaffold LLPS as much as strongly binding ligands. Furthermore, binding of ligands to non-sticker (spacer) regions rather than stickers of scaffolds is more favorable for enhancing LLPS (6, 7). Lastly, ligands with the same composition but different conformations or architecture can lead to highly different scaffold phase behaviors (8, 11–13). However, systematic studies are needed to determine the rules of how ligand architecture affects phase separation.

One prevalent set of ligands includes ubiquitin (Ub) and polyubiquitin (polyUb) chains, which are post-translationally attached to proteins and target them to different signaling outcomes, including proteasomal degradation and the DNA damage response, among many others (14). PolyUb chains, composed of different lengths and linkages, are a regulator of biomolecular condensates in protein quality control, autophagy, and immune system activation. Condensation can be driven by ubiquitination of E3 ligases such as TRIM3 (15), or regulated by ligase-mediated ubiquitination of proteins such as Disheveled-2 in Wnt signaling for organismal development (16). Cargo proteins modified with K63-linked polyUb chains interact with p62 to form pre-autophagosome condensates that later recruit autophagosome machinery (12, 17); furthermore, autophagy cargo adaptor NBR1 condenses with K63-linked and M1-linked chains (18). NEMO condenses with K63- and M1-linked polyUb chains, but not K6-, K11-, K29-, K33- and K48-linked polyUb chains, to facilitate activation of the IKK complex leading to downstream NF-κB processing (11, 19). In contrast, hHR23B preferably binds to and condenses with K48-linked chains to form nuclear condensates and recruit the proteasome and other protein quality control components to degrade defective ribosomal proteins (13). These different polyUb chains, which differ *only* in the location of the isopeptide bond that connects the Ub units, signal for different cellular outcomes (the Ub code) through the regulation of condensate formation. Therefore, these studies suggest that the differential effects on the condensation of interacting proteins is a mechanism for the Ub code to be read and interpreted in the cell (14, 20). Given the growing evidence for polyUb chain involvement in condensate formation, it is essential to determine the molecular rules by which polyUb chains regulate biomolecular condensates.

We recently showed that polyUb chain conformation contributes to whether polyUb chains promote or inhibit phase separation of UBQLN2, a Ub-binding shuttle protein critical to cellular protein quality control mechanisms. UBQLN2 phase separation is driven by homotypic interactions involving multiple folded and intrinsically-disordered regions across the protein (21, 22). MonoUb drives disassembly of UBQLN2 condensates (21). However, compact tetraUb (Ub4) chains (K11, K48) largely inhibit phase separation of UBQLN2, whereas extended tetraUb chains (K63, M1) promote phase separation of UBQLN2. Via binding interface mapping and K_d_ measurement, we showed that UBQLN2 interacts similarly with these different chain types (8, 23). These collective observations suggest that the distinction between linkages manifests through the emergent property of condensation. These studies led us to hypothesize that the more extended the polyUb chain, the greater the ability to promote UBQLN2 phase separation. To systematically examine the features of polyUb ligands that regulate phase separation, we employed a library of designed polyUb hubs where we modified either the linker between the Ub units on these hubs, or the binding affinity between the Ub unit and UBQLN2. By combining information on the solution structures of these designed hubs and changes in UBQLN2 phase diagrams with theoretical modeling, we established a set of rules regarding the recruitment of these ligands into UBQLN2 condensates. These rules, together with previously established ones, enable the prediction of how ligands modulate phase behavior and determine the designs of ligands for tighter control of specific condensate assembly as a function of ligand concentration.

## Results

### Decreased binding affinity between Ub and UBQLN2 weakens polyUb ability to promote UBQLN2 phase separation

We previously showed that Ub interacts with UBQLN2 at the same UBA residues that are involved in LLPS-driving multivalent interactions within UBQLN2 (21). These Ub:UBA interactions can either inhibit or further promote UBQLN2 LLPS, depending whether Ub is in monomeric or tetrameric (Ub4) states, respectively (8). To investigate how different Ub:UBA binding affinities affect UBQLN2 LLPS, we introduced single and double amino acid substitutions at select residues (I44 & V70) on the Ub hydrophobic binding patch (Fig. 1A) (24–26). Using NMR spectroscopy (Fig. S1), we determined that all of the substitutions reduced the binding affinity between Ub and UBQLN2 to different extents in the following order: Ub > Ub V70I > V70A > I44V > V70I/I44V > V70A/I44V > I44A (Table S1).

**Figure 1.**
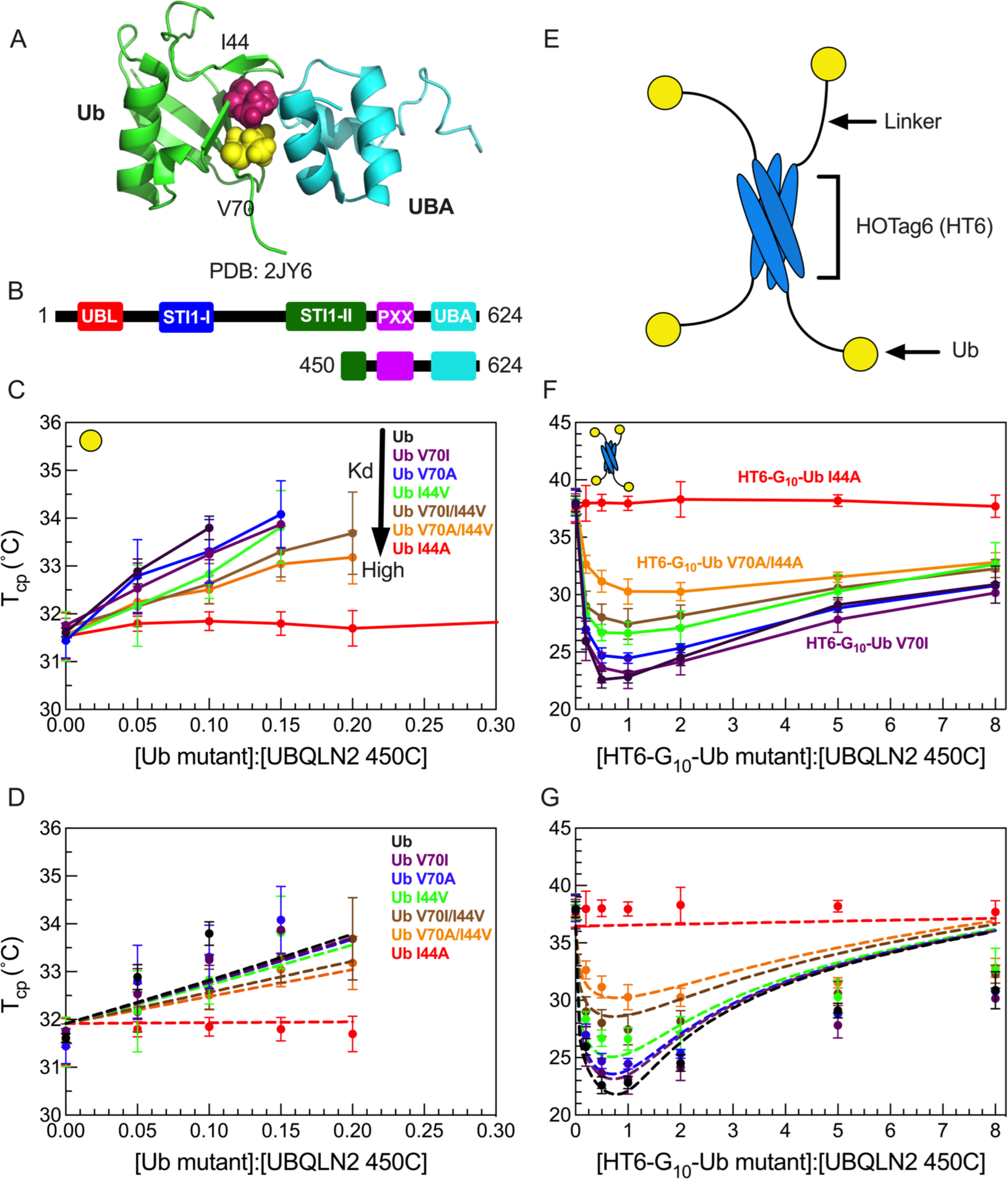
HT6-Ub variants with weakened binding affinity do not efficiently promote phase separation of UBQLN2/HT6-Ub. (A) Structure of Ub/UBA complex with the hydrophobic patch residues I44 and V70 of Ub represented in spheres. (B) Domain structure of full-length UBQLN2 and UBQLN2 450-624 construct (450C) used in this study. (C) Phase diagrams of 450C phase separation in the presence of monoUb variants. Phase separation occurs above phase boundary. (D) Comparison of theory (dotted lines) to experiments (points) for the monoUb variants in panel C. (E) Architecture of HOTag6-G_10_-Ub (HT6-Ub) ligand. (F) Phase diagrams of 450C phase separation in the presence of HT6-Ub variants. (G) Theory fit (dotted lines) to HT6-Ub variants using an inclusion energy of 9.5 kT. Data points and error bars in panels C and F reflect averages and standard deviations across at least n=6 experiments using two protein stocks.

We first quantified the effects of the substitutions in monoUb on the phase separation of a UBQLN2 C-terminal construct 450-624 (450C, Fig. 1B), which exhibits similar LLPS behavior to full-length UBQLN2 (21). We performed temperature-ramp turbidity assay experiments that monitor the change in A_600_ values (as a proxy for phase separation) at different Ub:450C ratios, but fixed 450C concentration and buffer composition. Using the cloud point temperature (T_cp_) obtained from these turbidity data, we built experimental temperature-component phase diagrams that delineate the coexistence curve, above which the UBQLN2:Ub solutions are phase-separated, and below which they are not. As the binding affinity between UBQLN2 and Ub variants decreased, Ub variants were less effective at binding to UBA and inhibiting 450C LLPS, which occurred at lower temperatures and higher Ub:450C ratios (Fig. 1C). The I44A substitution completely disrupted Ub:UBA binding, hence Ub I44A had little or no effect on 450C phase separation (Fig. 1C).

We next quantified the effects of the substitutions in the designed tetrameric HOTag6-(G)_10_-Ub (HT6-Ub, Fig. 1E) on 450C LLPS (8). HT6-Ub, a structural mimic of a multi-monoubiquitinated substrate, can significantly promote UBQLN2 phase separation over a wide range of HT6-Ub:UBQLN2 ratios (8) (Fig. 1F). Temperature-composition phase diagrams showed that as the binding affinity between UBQLN2 and HT6-Ub variants decreased, HT6-Ub variants were less effective at enhancing 450C LLPS (Fig. 1F). Together, data for both Ub and HT6-Ub variants demonstrated that binding affinities are positively correlated with the ability of the ligand Ub/HT6-Ub to modulate the phase behavior of the LLPS-driver 450C. These trends are consistent with results from a previous theoretical study using a generic LLPS-driver and its mono- and di-valent ligands (6).

### Theory development for how Ub ligands inhibit and promote phase separation

We developed a theory to reconcile the observation that monoUb and polyUb inhibit and promote phase separation, respectively. The key elements of the theory (see SI) include the enforcements of the chemical equilibrium of polyUb-UBQLN2 binding and the equilibrium of molecular partitioning between phases (dilute/dense). Here, we consider UBQLN2 as a phase separation “driver”, which is a macromolecule capable of phase separation on its own. The Ub or polyUb ligand is termed a “hub”, by which we refer to the ligand’s ability to interact with multiple UBQLN2 molecules, depending on the number of Ub units present on the hub. We assume that a driver has a single binding site for the hub, in accord with our experimental data where UBQLN2 binds to a single Ub molecule. Chemical equilibrium is enforced via the equilibrium constants for the reaction: *h + n d ⇌ hd_n_*:

(Eq. 1)

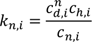

where *k*_*n*,i_ is the dissociation constant for a driver/hub complex (oligomer) containing one hub molecule and n drivers (henceforth referred to as an n-mer). *c*_*d*_, *c_h_*, and *c*_*n*_ are the concentrations of monomer (unbound) drivers, monomer (unbound) hubs, and n-mers, respectively, and the i-index indicates either the dilute phase (V = vapor) or the dense phase (L = liquid). Partitioning equilibrium is enforced by introducing transfer free energies *s_h_* and *s*_*d*_:

(Eq. 2a, 2b)

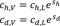

where *s_h_* > 0 indicates unfavorable transfer for the hub to the dense phase (the hub is unable to phase separate independently) and *s*_*d*_ < 0 indicates favorable transfer for the driver to the dense phase. The transfer energies, and all other energies, are given in units of k_B_T, where T is the absolute temperature and k_B_ is the Boltzmann constant. The favorable interactions captured by *s*_*d*_ will be temperature-dependent, which can be incorporated in the theory by noting that in the absence of hub molecules

(Eq. 3)

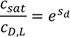

where *c*_*sat*_ is the saturation concentration of the driver at a particular temperature (see SI) and *c*_*D*,*L*_is the total concentration of driver in the liquid (henceforth capital H and D indices indicate total concentrations of hubs and drivers while lowercase h and d indicate concentration of hubs and drivers in the unbound state). Combining Eq. 2 and Eq. 3, we find

(Eq. 4)

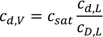

This equation says that phase separation can occur with *c*_*d*,*V*_ below the *c*_*sat*_ if the concentration of unbound driver in the dense phase is less than the concentration of driver in the homogenous driver fluid (UBQLN2 phase separation alone). This condition is easy to attain in the presence of hubs because n-mers will occupy a significant fraction of the dense phase volume such that 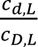 <1. In other words, n-mers dilute the concentration of unbound drivers in the dense state, which lowers their chemical potential and allows phase separation to occur at lower concentration. To determine a condition for the onset of phase separation, we use the fact that when polymers phase separate the mass concentration in the dense phase is insensitive to the molecular weight (or equivalently, polymerization number) of the individual molecules (5). This is because the microscopic interactions and mesh structure of the dense phase are both much smaller than the molecules. In agreement with this expectation, the concentration of UBQLN2 molecules in the dense phase is nearly unchanged as polyUb is added (8). Therefore, our criterion for phase separation is that the total concentration of drivers in the dense phase is equal to the concentration of the homogeneous fluid

(Eq. 5)

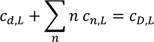

With the use of Eqs 1-4, this condition can be cast in terms of the dilute phase concentrations (see SI).

(Eq. 6)

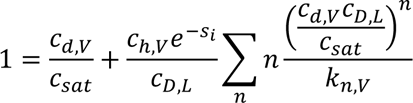

We refer to the quantity *s*_i_ = *s_h_* + Δ*s* as the “inclusion energy”. It accounts for two detrimental effects from transferring an n-mer to the dense phase. The first, described by *s_h_*, is the disruptive effect of inserting the hub within the network of attractive interactions formed by the drivers. In our model, this contribution depends only on the number of Ub modules in a hub. The second effect, captured by Δ*s*, comes from the fact that drivers bound to the hub are constrained in their interactions with the surrounding fluid so that the driving force for an n-mer to enter the dense state may differ from *n* ∗ *s*_*d*_. We expect that these constraints will depend on the specific linkage and/or geometry of the hub. The total inclusion energy, *s*_i_, is the only free parameter in Eq. 6.

### HT6-Ub imparts a significant inclusion energy that is offset by Ub binding affinity to promote UBQLN2 phase separation

We tested our theoretical model on our 450C data for monoUb and HT6-Ub variants (Fig. 1D and 1G). First, we performed a global fit to the experimental phase diagrams for HT6-Ub variants of different Ub:UBA binding affinities. The global fit was performed using the experimentally obtained binding affinities between monoUb and UBA (Fig. S1, Table S1). We assumed that each Ub in HT6-Ub interacted with the UBA of a single 450C with the same K_d_ as the isolated Ub:UBA interaction; previous data suggested this is the case for HT6-Ub and UBQLN2 (8). Thus, a HT6-Ub molecule can maximally have four 450C molecules bound. Importantly, the theory-derived coexistence curves mirrored the experimental phase diagrams for HT6-Ub variants of different Ub:UBA binding affinities (Fig. 1G). Stronger binding between Ub and UBA stabilized HT6-Ub:450C phase separation at low Ub:UBQLN2 ratios with the lowest T_cp_ generally near 1:1 stoichiometry of Ub:450C. The best fit value of the inclusion energy was 9.5 kT, suggesting that it costs almost 2.5 kT to transfer each ubiquitin unit to the dense phase (Fig. 1G). This value is very close to the energy of inserting RFP tags into SUMO-SIM condensates (27). The Ub inclusion penalty is offset by the favorable binding between the multiple Ub units on the hub and the 450C molecules. Thus, the reentrant behavior at stoichiometries greater than 1:1 is due to hubs that incur the full 9.5 kT inclusion energy but, having less than four drivers bound, have a reduced driving force for phase separation. We next considered the monoUb ligand. Using the same parameters, and an inclusion energy that is one-fourth of the HT6-Ub inclusion energy, we also recapitulated the phase diagrams for the monoUb variants (Fig. 1D). These results suggest that a single Ub unit is unfavorable to enter the dense phase, thus driving destabilization of UBQLN2 phase separation.

We note that the theory curves do not capture the phase boundary at higher Ub:UBQLN2 ratios; the theory predicts less phase separation (higher T_cp_) than what we observe (lower T_cp_). We believe this discrepancy is due to cooperativity in the binding of drivers to the hub. We speculate that the same favorable driver-driver interactions that promote phase separation will make it more likely for drivers to bind to hubs that already have drivers bound. Thus, when there is insufficient amount of drivers to saturate the hubs, the hubs will prefer fully-bound or fully-empty states, rather than the randomly occupied states assumed by the theory. Because of the unfavorable hub inclusion energy, a smaller number of fully bound hubs is more favorable to phase separation than a larger number of partially bound hubs. Due to the systematic overestimation of the phase boundary at high stoichiometries, we only used Ub:UBQLN2 ratios less than or equal to 2.0 to fit the model parameters.

### HT6-Ub with longer linkers are more extended in solution and less effective at enhancing UBQLN2 phase separation

Our recent work demonstrated that more extended polyUb chains promoted phase separation of UBQLN2, with the most extended HT6-G_10_-Ub and the most compact K48-linked Ub4 exhibiting the most and least LLPS-enhancing ability, respectively (8). However, HT6-G_10_-Ub and polyUb chains of different linkages differ not only in conformation, but also in overall topology, steric hindrance within the polyUb chains, accessibility to the Ub hydrophobic binding surface, and orientation of the Ub units, stymieing clear interpretation of the observed correlation between ligand conformations and ligand ability to modulate LLPS. Moreover, we hypothesize that there is a limit to the positive correlation between the extendedness of polyUb hubs and the enhancement of UBQLN2 phase separation. Beyond this limit, the Ub units in more extended polyUb hubs would be too far apart to bring the driver molecules close together to enhance phase separation. Here, we aimed to systematically investigate how changes in polyUb hub conformation modulate UBQLN2 LLPS by using the simplicity of our designed tetrameric HT6-Ub, the architecture of which enabled accommodations for linkers of varying lengths and flexibility without interfering with the Ub units. We created a library of linkers comprising multiple GS (Gly-Ser) or PA (Pro-Ala) blocks (Fig. 2A, Fig. S2), which are flexible and rigid in conformation, respectively (28).

**Figure 2.**
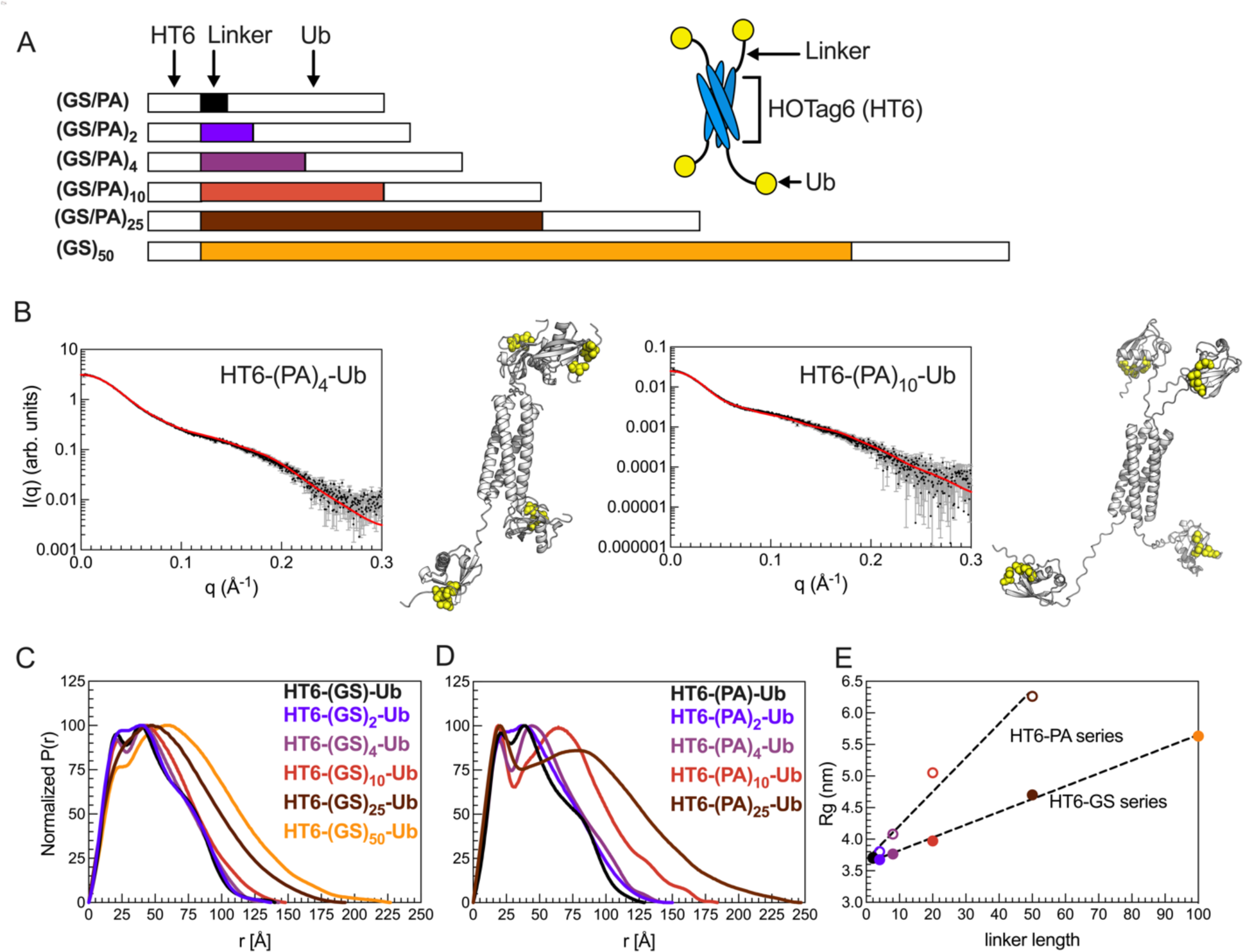
HT6-Ub constructs with longer linkers are larger in size and more extended in overall conformation. (A) Library of HT6-Ub constructs with different linkers with a cartoon representation of the spider-like HT6-Ub geometric architecture. (B) Experimental X-ray scattering curves (black) for designed HT6-(PA)_x_-Ub constructs overlaid with scattering curve predicted (red) from single best structure (shown) derived from SASSIE conformational ensembles. Hydrophobic patch residues L8, I44, and V70 are represented as yellow spheres. (C,D) P(r) profiles from SAXS experiments on HT6-Ub ligands of different linker lengths. (E) Radius of gyration (R_g_) as determined from Guinier fits to SAXS experiments for HT6-Ub constructs (closed circles: GS series and open circles: PA series) as a function of linker length (# of amino acids). Black dotted lines represent simple linear regression of each data set.

We performed size-exclusion chromatography coupled with multiangle light scattering and small-angle X-ray scattering (SEC-MALS-SAXS) on the HT6-Ub constructs with different GS and PA linkers to gain insights on how these linkers affected the overall conformations and structures of the HT6-Ub hubs. SEC-MALS experiments confirmed that each HT6-Ub construct examined herein existed as a tetramer in solution and that elution time, hence overall size of the HT6-Ub complex, increased with linker length (Fig. S3, Table S2). Representative structures of these HT6-Ub constructs were determined by refining AlphaFold-predicted starting structures (29) against our scattering data using conformational ensembles constructed with SASSIE (30) (Fig. 2B). Pair distance distribution function (P(r)) analysis showed that the HT6-Ub constructs with longer linkers adopt more extended structures (Fig. 2C). Guinier analysis of collected SAXS data also displayed a positive linear relationship between linker length and radius of gyration (R_g_), with R_g_ for constructs with (PA)_x_ linkers increasing more than with (GS)_x_ linkers as a function of linker length (Fig. 2D, Fig. S4, Table S2). HT6-Ub with PA linkers were much more extended than those containing GS linkers of the same length, consistent with (PA)_x_ being more rigid than (GS)_x_. For all HT6-Ub constructs used, the Ub units do not exhibit significant changes to structure or dynamics compared to monoUb, as observed by NMR spectroscopy experiments (Fig. S5A-E).

To compare flexibility of Ub tethered to HT6 to that of monoUb on a residue-by-residue basis, we performed NMR spin relaxation experiments on backbone amide ^15^N resonances. ^15^N R_1_ & R_2_ relaxation rates monitor backbone dynamics on millisecond to nanosecond timescales. MonoUb has the highest R_1_ rate of 1.73 s^-1^while Ub tethered to HT6 by (PA)_2_ and (GS)_2_ linkers showed the lowest R_1_ rate of 0.83 s^-1^ (Fig. S5). ^15^N R_1_ rates for Ub decreased while ^15^N R_2_ rates increased in the order of decreasing linker length: (Ub > HT6(GS)_50_Ub > HT6(PA)_10_Ub > HT6(GS)_10_Ub > HT6(G)_10_Ub > HT6(PA)_2_Ub ∼ HT6(GS)_2_Ub). Decreases in R_1_ rate with corresponding increases in R_2_ rate indicate an overall slower rotational tumbling rate for the protein; hence Ub tethered to HT6 with a shorter linker tumbled more slowly than Ub tethered with a longer linker. Collective observations from spin-relaxation and SEC-MALS-SAXS experiments indicated that Ub tethered to HT6 with a longer linker exhibited a faster tumbling rate (has a higher degree of freedom to move) and adopted more extended structures.

We next performed turbidity assays of 450C in the presence of HT6-Ub constructs containing different linkers. The resulting phase diagrams significantly shifted as a function of linker length (Fig. 3). Importantly, the overall shape of these phase diagrams (minima and slopes with increasing Ub:450C ratios) remained unchanged as linker length was modulated, unlike that of the binding variant HT6-Ub/450C phase diagrams (Fig. 1F), indicating that the Ub:450C binding surface was unperturbed. Indeed, NMR titration experiments reported similar *K*_d_ values and chemical shift perturbations (CSPs) between UBA and the Ub units across several HT6-Ub constructs (Table S1, Fig. S6). These results are consistent with those from our prior work which showed that differences in UBQLN2 phase separation behavior do not correlate with changes in *K*_d_ between polyUb of different linkages and UBQLN2 (8). Therefore, these results suggest that the differences in the phase diagrams for HT6-(GS/PA)_x_-Ub and 450C stemmed solely from changes to the HT6-Ub linkers.

**Figure 3.**
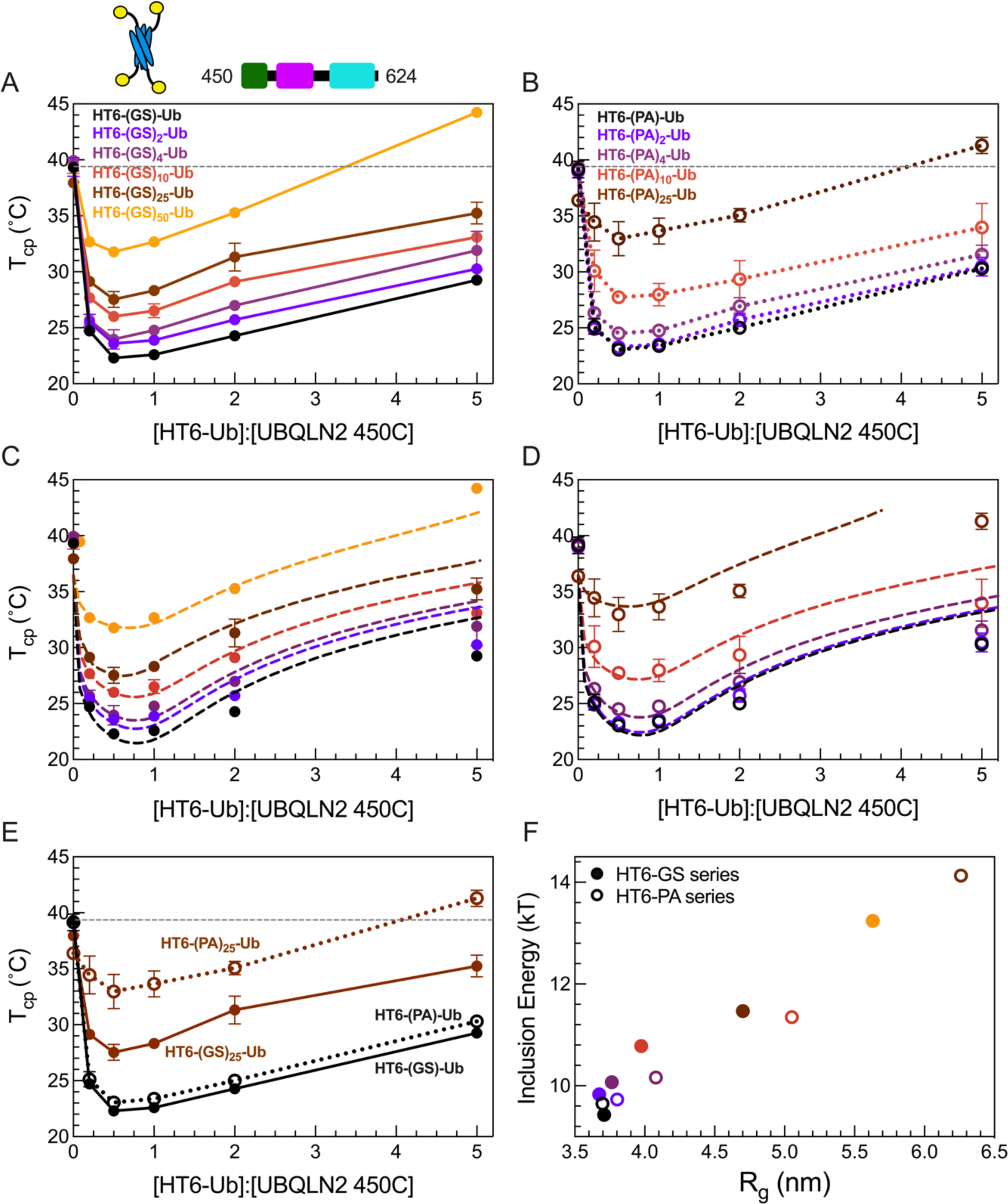
HT6-Ub constructs with longer linkers are less effective at enhancing phase separation of UBQLN2. (A,B) Phase diagrams of 450C phase separation in the presence of HT6-Ub variants containing variable GS linker length (A) or variable PA linker length (B). Data points and error bars in panels A and B reflect averages and standard deviations across at least n=6 experiments using two protein stocks. Cartoon above (A) denotes use of HT6-Ub ligands with 450C (fixed concentration of 50 µM) for these experiments. Phase separation of 450C and HT6-Ub occurs at temperatures above (but not below) phase boundaries. (C,D) Theory fit (dotted lines) to HT6-variants. (E) Comparison of 450C phase diagrams with HT6-GS-Ub (filled circles, black), HT6-PA-Ub (open circles, black), HT6-(GS)_25_-Ub (filled circles, brown), and HT6-(PA)_25_-Ub (open circles, brown). (F) Inclusion energy (kT) as determined from theory compared to the R_g_ of each HT6-Ub ligand used (as color-coded in panels C and D).

Our results show that the HT6-Ub constructs with the shortest linkers (HT6-GS-Ub & HT6-PA-Ub) were the most effective at enhancing 450C phase separation (lower T_cp_), and increasing either GS or PA repeat length diminished the ability of HT6-Ub to promote 450C phase separation over a wide Ub:450C range (Fig. 3A and 3B). Indeed, HT6-Ub constructs with longer (PA)_x_ linkers, which are much more extended than those with longer (GS)_x_ linkers of the same length (Fig. 3E), were less effective at enhancing 450C LLPS compared to their (GS)_x_ counterparts. These results appear to contradict our previous finding that more extended polyUb chains can more effectively enhance UBQLN2 LLPS (8). To understand this contradiction, we used our theoretical model to determine the inclusion energy. We assume that these inclusion energies are the sum of the linkage-independent effect of inserting the hubs into the condensate (s_h_) and a linkage-dependent effect on driver-driver interactions (ΔS) (see SI). This analysis revealed that the inclusion energy of introducing the HT6-Ub hub into the dense phase was lowest (most favorable) for hubs with the shortest linker and increased for hubs with longer linkers (Fig 3B, D, F). Together, our experimental and modeling results suggest that the HT6-Ub hubs with shorter linkers are better accommodated within the condensate of drivers, hence lowering the inclusion energy for hub incorporation and enhancing LLPS. Our data suggest that the HT6-Ub constructs with the shortest linkers are already optimized to maximize phase separation with UBQLN2. Our previous work (8) included more compact chains (K48-, K63- and M1-Ub4) and a more extended HT6-Ub construct (with a 10-residue linker), but not the most optimal HT6-Ub constructs (with a two-residue linker) studied here.

### Designed linear(M1-linked) polyUb chains with longer linkers are less effective at enhancing UBQLN2 phase separation

Ligand hub architecture could affect phase separation of drivers (e.g. UBQLN2) via differences in hub geometry, binding constraints, and the proximity of bound drivers. The HT6-Ub hub comprises Ub units that can rotate independently from one another. In contrast, the Ub units in polyUb chains are covalently attached to one another and thus experience more constraints. Therefore, the trends observed for HT6-Ub with varying linker lengths might differ from those observed for polyUb chains with varying linker lengths. Since conjugation of polyUb chains requires specific recognition by particular Ub-conjugating enzymes, most chains are not amenable to manipulation. One exception is the linear M1-linked chain in which the C-terminus of one Ub is covalently linked to the N-terminal amine of the next, and so forth; this chain type adopts extended conformations in solution (8, 31). Moreover, structural and NMR dynamics studies (32, 33) revealed that the last four residues of Ub (residues 73-76) are unstructured and very flexible. Therefore, this linear architecture enabled us to create a large set of M1-Ub4 constructs that contained both shorter and longer linkers than the naturally-occurring M1-Ub4 to examine the effects of changing the spacing between Ub units on UBQLN2 phase separation (Fig. 4).

**Figure 4.**
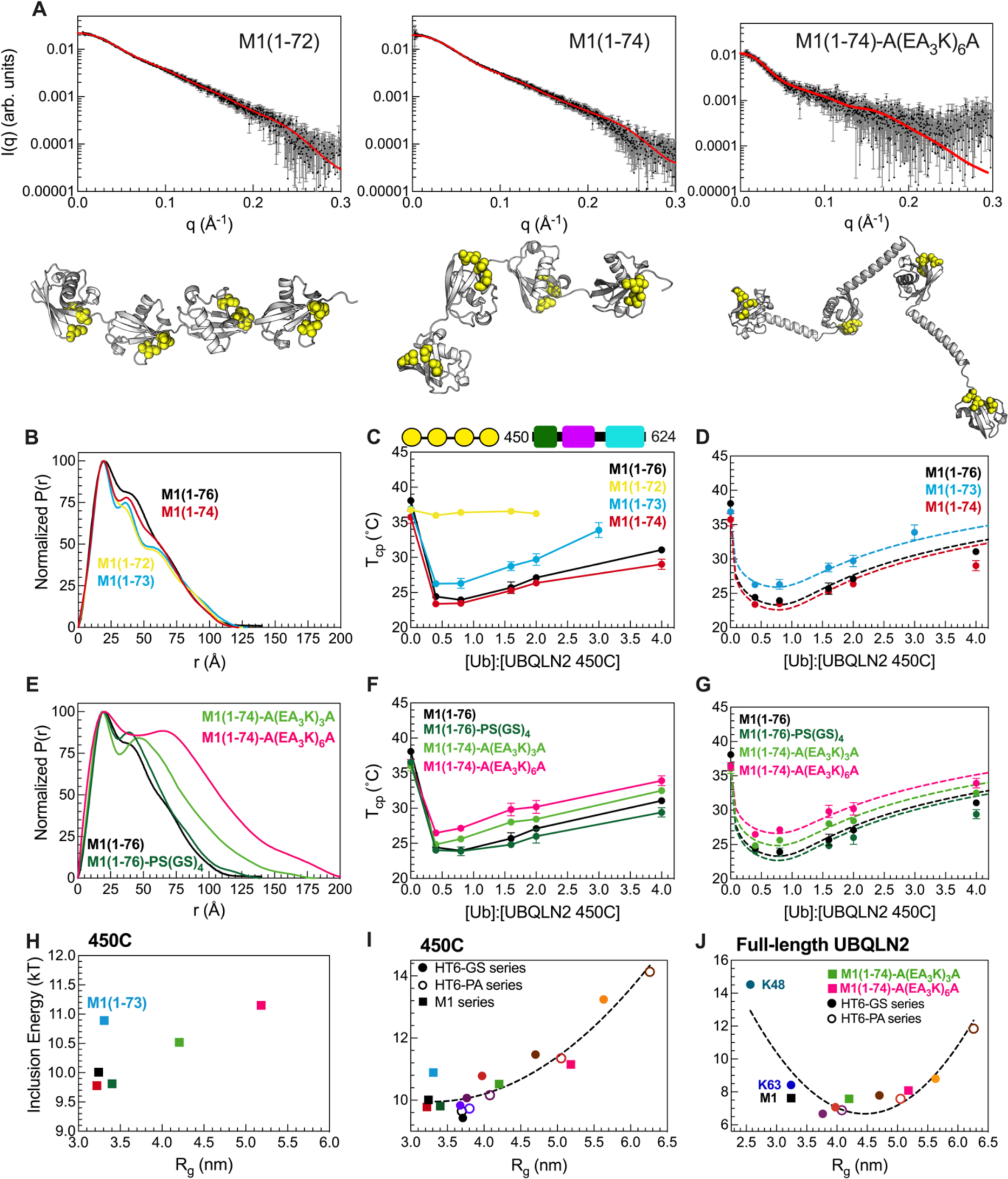
Designed linear polyUb chains with either short or long Ub-Ub linkers are less effective at enhancing phase separation of UBQLN2. (A) Experimental X-ray scattering curves (black) for designed M1-Ub4 constructs overlaid with scattering curve predicted (red) from single best structure (shown) derived from SASSIE conformational ensembles. Hydrophobic patch residues L8, I44, and V70 are represented as yellow spheres. (B) P(r) profiles of designed M1-Ub4 constructs with shorter Ub-Ub linkers. (C) Temperature-concentration phase diagrams of 450C with designed M1-Ub4 constructs. Phase separation of 450C and M1-Ub4 constructs occurs at temperatures above (but not below) phase boundaries. (D) Theory-derived curves overlaid on experimental data. (E) P(r) profiles of designed M1-Ub4 constructs with longer Ub-Ub linkers. (F) Temperature-concentration phase diagrams of 450C (fixed concentration of 50 µM) with designed M1-Ub4 constructs. (G) Theory-derived curves overlaid on experimental data. (H) Theory-derived hub inclusion energy for M1-Ub4 constructs of various Ub-Ub linker lengths (as color-coded in panels D and G). Label is added for M1-Ub4 (1–73) as this construct binds less effectively to 450C. (I) For 450C phase separation, theory-derived hub inclusion energy for all designed polyUb hubs in this study. Data points are color-coded as in panels D and G, and from Fig. 3F. Inclusion energies for M1-Ub4 and HT6-Ub constructs are represented as squares and circles, respectively. (J) For full-length UBQLN2 phase separation, theory-derived hub inclusion energy for naturally-occurring Ub4 polyUb chains, two designed M1-Ub4 constructs of longer Ub-Ub linker lengths, HT6-(GS)_x_-Ub hubs (filled circles), and HT6-(PA)_x_-Ub hubs (open circles). Parabolic trendlines (dotted lines) in panels I and J are drawn to guide the eye. R_g_ values in panels H, I, and J are of the different ligand hubs used.

First, we examined shorter linkers, i.e. by deleting residues from the C-terminus of Ub for all three Ub-Ub linkers in a M1-Ub4 construct ((1-72, 1-73, 1-74, and 1-76 (WT)). Although the scattering profiles and R_g_ values of these different constructs were relatively unchanged (Fig. 4A, Fig. S7), distinct peaks appeared in the M1-Ub4 (1–72) and M1-Ub4 (1–73) P(r) profiles (Fig. 4B). We next compared SASSIE-constructed conformational ensembles for each construct against the experimental SAXS scattering curves (Fig. 4A). This analysis revealed that the Ub units in M1-Ub4 (1–72) exhibited restricted mobility due to the short linkers between Ub units. NMR spectra of these chains were similar to that of monoUb, indicating that the Ub units remained well-structured (Fig. S8). However, CSPs between M1-Ub4 (1–72) or M1-Ub4 (1–73) and monoUb suggested that the hydrophobic patch residues (8,44,70) made Ub-Ub contacts not present in naturally-occurring M1-Ub4 or the HT6-Ub hubs (Fig. S8A-D). NMR titration experiments revealed that binding between M1-Ub4 (1–72) and UBQLN2 UBA is significantly attenuated (Fig S8G). Modeling of the UBA to SASSIE structures of M1-Ub4(1–72) suggested that multiple UBA domains can’t be accommodated on this shortened Ub4. In line with our structural data, phase diagrams of 450C with M1-Ub4 (1–74) and M1-Ub4 (WT) were very similar and almost within error of each other (Fig. 4C,D). However, M1-Ub4 (1–73) was less effective at enhancing 450C phase separation than M1-Ub4 (1–74), and M1-Ub4 (1–72) had no effect on 450C phase separation. These results are in agreement with our previous data on the compact K48-Ub4 chains, the Ub units of which interact with one another and impede the ability of the chain to promote UBQLN2 LLPS.

Next, we examined M1-Ub4 constructs with increased linker length between Ub units (Fig. 4E-G). The overall SAXS scattering profile and R_g_ for an M1-Ub4 chain with a 10-residue linker (PS(GS)_4_) between each Ub unit were largely unchanged compared to M1-Ub4 (WT) despite the addition of these 10 residues (Fig. 4E). In line with these solution properties, the overall phase diagram of 450C with M1-Ub4 (PS(GS)_4_) is similar to M1-Ub4. We hypothesized that the (PS(GS)_4_) linker following G_75_G_76_ in each Ub is too flexible to elicit change in conformation of M1-Ub4. Therefore, we installed sequences A(EA_3_K)_3_A or A(EA_3_K)_6_A which are predicted to be helical (28) between the Ub units in a M1 (1–74) construct. SAXS data suggested that the structures of these M1-Ub4 constructs are significantly more extended compared to M1-Ub4 (1–76) as R_g_ values increased by nearly 10 Å (Table S2). Experimental phase diagrams of 450C with these polyUb hubs of longer linkers showed that these hubs were less effective at enhancing 450C phase separation (Fig. 4H). Using our theory to disentangle the architecture-dependent contributions to the inclusion energy, we found that the inclusion energy is minimized for hubs containing linkers of optimal lengths (Fig. 4I). Our theory is unable to determine the inclusion energy for M1-Ub4 (1–72) as the hub is unable to bind to 450C molecules thus no longer meeting a critical assumption of our model (Fig. S8G). Furthermore, our data suggest that wild-type M1 (1–76) polyUb chains are well-optimized to enhance 450C phase separation at low Ub:450C concentrations, compared to other M1 polyUb chains where the linker is shortened or lengthened (Fig. 4).

Our data from the structurally distinct sets of M1-Ub4 and HT6-Ub constructs establish a general trend that compact hubs with unhindered binding ability are favored for inducing 450C phase separation (Fig. 4I, Table S4). The inclusion energies for the M1-Ub4 and HT6-Ub constructs nearly overlap when plotted as a function of R_g_. From these results, we can conclude that the geometry of the ligand hub is not as important as density/spacing of Ub units.

To determine if full-length UBQLN2 responds to polyUb hubs in a similar manner as the shorter UBQLN2 450C construct, we obtained UBQLN2 phase diagrams in the presence of linear M1 polyUb chains with longer linkers (A(EA_3_K)_3_A and A(EA_3_K)_6_A) as well as HT6-Ub hubs with different GS or PA linker lengths, and combined these results with recent work on polyUb of different linkages (K48, K63, M1) (8) (Fig. S9). Consistent with our 450C data above, hubs with linkers that are either too short or too long (have higher inclusion energy) do not enhance UBQLN2 phase separation compared to hubs with optimal linkers (e.g. HT6-(GS)_4_-Ub) (Fig. 4J, Table S3). Our results reveal that the free energy cost of inserting hubs into the dense phase is dependent on the spacing of Ub binding sites. We propose that this Ub-Ub spacing is part of the Ub code that regulates phase separation via polyUb chains of different lengths and linkages, as well as by ubiquitination at multiple sites of target proteins (represented by the HT6-Ub constructs in this study). The different linkages used in the Ub code deliver biochemical information via Ub-Ub spacing.

## Discussion

The formation and disassembly of biomolecular condensates can be extensively tuned by ligand-mediated phase transitions (5, 34). As many ligands in the cell are multivalent by design, it is essential to establish the molecular rules that dictate how these ligand hubs can either inhibit or promote phase separation (35). Using a combination of experiment and analytical theory, we determined that, aside from previously examined properties, two other properties of the ligand hub contribute significantly: 1) effective linker length between binding sites within the hub and 2) binding affinity between the hub and phase-separating driver molecules. We find that the specific geometry of the ligand hub matters less than the binding site spacing in hubs. These tunable features of multivalent ligand hubs modulate multicomponent phase separation as demonstrated for the polyUb-UBQLN2 system.

The analytical theory developed here is a framework to consider how ligand hubs impact phase separation of driver molecules. For ligands to be incorporated into the dense phase of driver condensates, there is an energetic penalty as a consequence of the ligand disrupting the driver-only interactions in the dense phase. We show that this energetic penalty is tuned by the physicochemical properties of the ligand hub, including valency, binding site spacing, and binding affinity. Despite the energetic penalty, introducing multivalent polyUb ligand hubs into UBQLN2 condensates still promotes phase separation of UBQLN2 relative to UBQLN2 in the absence of a polyUb hub ligand. This is because the multivalency of the polyUb hub enables the accumulation of multiple sets of favorable binding energies between each Ub unit and UBQLN2. However, when spacing of the Ub units are too close or too far, polyUb hubs and UBQLN2 molecules are not in a favorable arrangement, thus destabilizing phase separation (Fig. 5). This argument suggests that non-ideal polyUb hubs (i.e. hubs with non-optimal Ub-Ub spacing) may introduce voids and/or steric clashes among the UBQLN2 molecules such that they are not in their ideal arrangement for both a) interaction with the polyUb ligand, and b) maintaining the condensate state.

**Figure 5.**
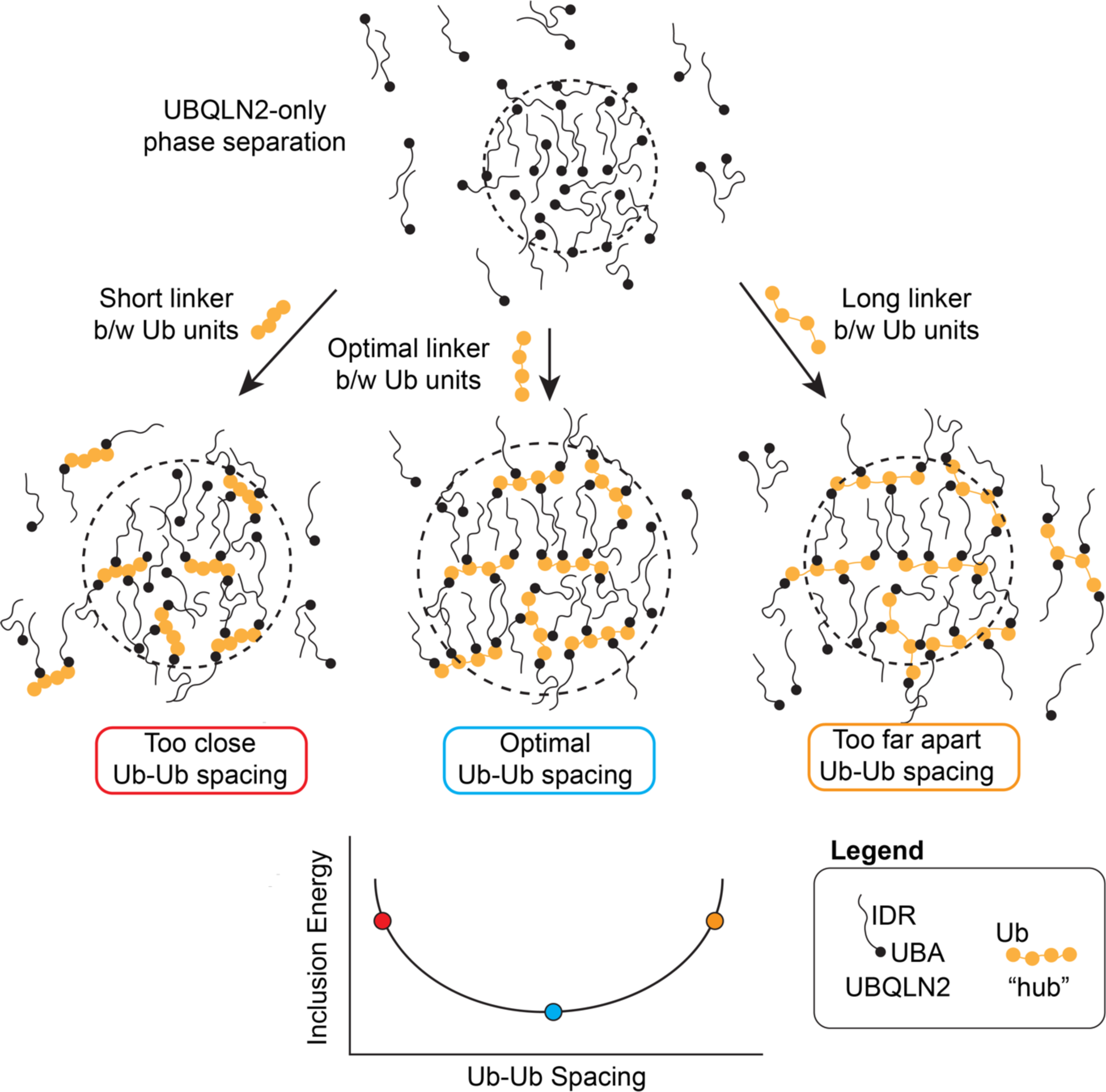
Introducing a ligand into condensates incurs an energetic penalty (inclusion energy) that can be reduced by optimizing the ligand architecture. Adding a ligand to the condensate disrupts the optimal binding of the surrounding driver (UBQLN2) molecules. This disruption can be minimized if the binding sites on the ligand are commensurate with the spacing of the surroundings. This optimal spacing minimizes the inclusion energy. In the presence of any ligand hub, phase separation of driver is enhanced (depicted here as a larger droplet size) and less UBQLN2 is found in the dilute phase. This represents the experimental observation that all of these multivalent hubs still promote phase separation of the driver molecules.

Here, we used two sets of designed polyUb hubs (HT6-Ub and M1-Ub4 chains of different Ub-Ub spacings) to dissect hierarchical contributions (36) to phase separation of driver/hub systems. We have not perturbed UBQLN2 (driver) molecules but focus exclusively on the effects of the ligand/hub. Despite the very different architecture of how the Ub units are arranged on the hubs (spider-like vs. beads on a string for HT6-Ub and M1-Ub4, respectively), several themes emerged from data with both types of hubs. First, this work and our prior report (8) showed that crowding the Ub units too close together disfavors UBQLN2 phase separation. Second, we showed here that increased distance between Ub units also disfavors UBQLN2 phase separation (Fig. 4). Third, because of these competing trends, there is an optimal spacing of Ub units that maximizes the propensity for phase separation (Fig. 4). Fourth, the optimal ligand hub size and Ub-Ub spacing is similar for both full-length UBQLN2 and 450C constructs (Fig. 4I and 4J). These data demonstrate the versatility of different ligand hub designs on effecting similar outcomes on phase separation of a scaffold like UBQLN2. Specifically, a multi-mono ubiquitinated substrate (e.g. HT6-Ub) could take the place of a polyubiquitin chain and promote phase separation of UBQLN2 over a wide range of Ub:UBQLN2 ratios.

Conceptually, our work shows that ligand hubs provide a sensitive mechanism to regulate condensate formation: 1) small differences in spacing between binding sites (e.g. Ub-Ub) can be magnified through driver phase separation (e.g. UBQLN2), and 2) phase separation of a driver can be driven to lower concentration with a ligand hub. Similarly, recent work on Speckle-type POZ protein (SPOP), a tumor suppressor that self-oligomerizes, has shown that SPOP cancer mutations can potentiate phase separation of SPOP with the DAXX substrate (37). DAXX has a low intrinsic tendency to phase separate, but SPOP can act as a “hub” that brings multiple DAXX molecules together to promote DAXX phase separation (38, 39). Wild-type SPOP requires a high DAXX:SPOP ratio to form the liquid-like condensates associated with DAXX ubiquitination. In contrast, cancer-associated SPOP mutations reduce SPOP oligomerization leading to more favorable arrangements between SPOP and DAXX to promote liquid-like SPOP/DAXX condensates at lower concentrations than for wild-type SPOP. In other words, cancer mutations optimize the SPOP “hub” to provide conditions that optimally promote DAXX phase separation. In a similar way, the polyUb hub optimizes UBQLN2 interactions to favor phase separation, but less optimal spacing between Ub units on the hub will disfavor UBQLN2 phase separation.

Interestingly, the Ub-Ub spacing in naturally-occurring M1-linked polyUb chains is generally optimal at promoting phase separation of UBQLN2 (Fig. 4C, 4F, Fig. S9). Shortening or lengthening the linkers by the removal or addition of multiple residues on the Ub C-terminal tail (Ub residues 73-76) reduced the ability of the polyUb hub to maximally promote phase separation of UBQLN2. This insight is important given that M1-linked polyUb chains are also best poised to drive multicomponent phase separation of other Ub-binding proteins such as NEMO and p62 (11, 12, 17, 19). These results suggest that the Ub code takes advantage of the optimal Ub-Ub spacing and the sensitivity of the high density (short spacer) side of the inclusion energy curve (Fig. 4I, J) in regulating condensate formation. The spacing within M1-linked polyUb chains are optimized for Ub-binding domains of highly diverse architectures, specifically a compact three-helix UBA domain located at one end of p62 and UBQLN2 in contrast to a long coiled-coil region in NEMO (40, 41). Moreover, M1-linked polyUb chains are optimized to both enhance the phase separation of systems that can phase separate on their own (UBQLN2) and drive the phase separation of systems that do not phase separate on their own (p62, NEMO). The heterotypic interactions lower the intrinsic c_sat_ (saturation concentration) of the Ub-binding partner, permitting regulation of condensate assembly/disassembly (5, 42). These data suggest that the architecture of M1-linked polyUb chains have favorable properties (flexibility and spacing of Ub binding sites) to promote phase separation of systems driven by either homotypic or heterotypic interactions.

Aside from Ub, other Ub-like modifiers of proteins, such as SUMO, UFM1, and NEDD8 form chains of different linkages or hybrid chains (i.e. Ub-SUMO or Ub-NEDD8 chains) (43, 44). SUMO and UFM1 contain unstructured regions of different lengths on either or both of their termini. Subsequently, the spacings between SUMO units on a chain are larger than those between Ub units. We showed that the UBA domain of UBQLN2 is unable to interact with M1-Ub4 (1–72) or M1-Ub4 (1–73) due to insufficient spacing between the Ub units. Just like the naturally-occurring M1-Ub4 (1–76) is optimized to accommodate several UBQLN2 molecules and enhance LLPS, the spacings between the units of SUMO chains, depending on linkage types, might have been optimized for binding to their partners (e.g proteins with multiple SIM modules). Protein polySUMOylation has been shown to regulate the assembly and dissolution of various biomolecular condensates, such as PML bodies, nucleoli, and stress granules (45). These multivalent Ub, UFM1, or SUMO chains could communicate their cellular code through modulating condensate assembly/disassembly. Therefore, knowledge of how changes in the spacing between Ub units affect UBQLN2 phase separation is also relevant to these Ub-like modifiers and their respective binding partners. This framework can be further extended to quantify the effects of other post-translational modifications on phase-separating systems (46, 47).

In summary, ligand hubs offer a tunable knob for modulating condensate assembly and disassembly. The interplay between ligand hub architecture and driver molecule arrangement leverages the sensitivity of phase separation in amplifying small differences, such as the distance between binding sites on a ligand hub. As a result, the total effect on condensates cannot be considered a sum of individual parts (e.g. specific interactions including driver-driver, ligand-driver) but rather an emergent property of the condensates (35). We expect that polyUb chains (homotypic linkages, mixed linkages, branched) and other polyUb-like systems (e.g. polyPARylation) use their multivalent architecture to dynamically regulate condensate properties and functions. For Ub and Ub-like proteins, we believe that the effect of these ligand hubs are the mechanistic underpinnings of the Ub code.

## Acknowledgements

This work was supported by NIH R01GM136946 (all protein purifications, turbidity assays, structural modeling, and NMR experiments) to C.A.C. and NIH R01GM141235 (theoretical modeling) to J.D.S. NMR data were acquired on an 800 MHz NMR spectrometer funded by NIH-shared instrumentation grant 1S10OD012254. This research used resources of the Advanced Photon Source, a U.S. Department of Energy (DOE) Office of Science User Facility operated for the DOE Office of Science by Argonne National Laboratory under Contract No. DE-AC02-06CH11357. This project was supported by grant P30 GM138395 from the National Institute of General Medical Sciences of the National Institutes of Health. Use of the Pilatus 3 1 M detector was provided by grant 1S10OD018090-01 from NIGMS. We acknowledge support and assistance from Dr. Jesse Hopkins and Dr. Maxwell Watkins on collecting SEC-MALS-SAXS data at APS. Suzanne Enos and Antara Chaudhuri were supported by NSF REU grant CHE1950802. We acknowledge Yiran Yang for assistance in purifying some protein for this project. We thank Susan Krueger, Tanja Mittag, and the Condensate Colloquium series (CCS) for scientific discussions on this project.

## Materials and Methods

### Subcloning, Protein Expression, and Purification

Ub and Ub mutants were expressed in *Escherichia coli* NiCo21 (DE3) cells and purified using previously described procedure (48).

The genes encoding all HT6-Ub constructs were codon-optimized, synthesized, and cloned into pET24b (Novagen) by GenScript (NJ, USA). HT6-Ub constructs were grown in NiCo21 cells in Luria-Bertani (LB) broth with 50 mg/L kanamycin to OD_600_ of 0.6 (except for HT6(GS)_50_Ub, which was grown to OD_600_ of 0.8-1.0), induced with 0.5 mM IPTG and expressed overnight at 37°C. Then the cells were pelleted, frozen, and resuspended to lyse in 20 mL of 50 mM Tris buffer pH 8.0 with 1 mM PMSF, 1 mM MgCl_2_, 1 mg/ml lysozyme and 25 µM Pierce universal nuclease. The lysate was centrifuged at 20 000 *g for 30 min, the supernatant was separated and loaded onto anion exchange column (Cytiva) and eluted with a gradient between 20 mM HEPES, 0.02 % NaN_3_ buffer pH 7 and 20 mM HEPES, 0.02 % NaN_3_ and 1 M NaCl buffer pH 7. Fractions containing HT6-Ub were collected and diluted 1:1 using 50 mM ammonium acetate pH 4.5. The resulting solution was centrifuged at 10 000 *g for 10 min, after which the supernatant was loaded onto a cation exchange column (Cytiva) and eluted with a gradient (Buffer A: 50 mM ammonium acetate pH 4.5, Buffer B: 50 mM ammonium acetate, 1 M NaCl pH 4.5). Fractions containing HT6-Ub were collected and dialyzed into 20 mM sodium phosphate buffer pH 6.8 with 0.5 mM EDTA and 0.02 % NaN_3_ overnight. Finally, the fractions were concentrated using Amicon Ultra concentrators (10,000 MW cutoff) and stored at −80°C until needed.

All M1-Ub4 constructs were expressed in *Escherichia coli* NiCo21 (DE3) cells in LB broth at 37°C overnight. Then the cells were pelleted, frozen, and resuspended to lyse in 20 mL of 50 mM Tris buffer pH 8.0 with 1 mM PMSF, 1 mM MgCl_2_, 1 mg/ml lysozyme and 25 µM Pierce universal nuclease. The lysate was centrifuged at 20 000 *g for 30 min, the supernatant was separated, and loaded onto the anion exchange column and the flowthrough was collected. The flowthrough was loaded onto the cation exchange column and eluted with a gradient (Buffer A: 50 mM ammonium acetate pH 4.5, Buffer B: 50 mM ammonium acetate, 1 M NaCl pH 4.5). The fractions containing M1-Ub4 were collected, concentrated, and buffer exchanged into 20 mM sodium phosphate buffer pH 6.8 with 0.5 mM EDTA and 0.02% NaN_3_ using Amicon Ultra concentrators (10,000 MW cutoff) and stored at −80°C until needed.

Full-length UBQLN2 and UBQLN2 450-624 were expressed and purified as described previously (21, 49). Briefly, the constructs were transformed into Rosetta 2 (DE3) pLysS *E. coli* cells and were grown in LB media with a final concentration of 50 mg/L kanamycin and 34 mg/L chloramphenicol to OD_600_ of 0.6-0.8, induced with 0.5 mM IPTG, and expressed overnight at 37°C. Then the cells were pelleted, frozen, lysed, then purified via a “salting out” process, where NaCl was added to the spun down lysate to a final concentration of 0.5 - 1 M. The resulting UBQLN2 condensate phase was pelleted and then resuspended in cold 20 mM sodium phosphate buffer pH 6.8, with 0.5 mM EDTA & 0.02 % NaN_3_. Leftover NaCl was removed through HiTrap desalting column (GE Healthcare), concentrated, and stored at −80 °C until needed.

### Spectrophotometric Absorbance/Turbidity Measurements

Turbidity assays were generally performed as described in (49). Protein samples were prepared by mixing different ratios of UBQLN2 450C (or full length UBQLN2) and Ub mutants/HT6-Ub/M1-Ub4 constructs (the initial concentrations of the protein stocks were doubled compared to the sample concentrations) in cold 20 mM sodium phosphate buffer pH 6.8 with 0.5 mM EDTA and 0.02 % NaN_3_. Then, cold 400 mM NaCl in 20 mM sodium phosphate buffer pH 6.8 with 0.5 mM EDTA and 0.02 % NaN_3_ was added in a 1:1 ratio. Both solutions were kept on ice for at least 5 min before mixing. Absorbance at 600 nm was monitored as a function of temperature using a Cary 3500 UV/Vis spectrophotometer (Agilent) using a temperature ramp rate of 1 °C/min increasing from 15 °C to 60 °C (for HT6-Ub, HT6-Ub mutants & M1-Ub4 constructs) and 25 °C to 60 °C (for Ub mutants). Several full-length UBQLN2 assays used a temperature range from 4 °C to 60 °C. Net absorbance values were recorded after subtracting the absorbance value from the reference buffer.

### Phase Diagram Measurements

The phase boundary for UBQLN2 LCST (lower critical solution temperature) phase transitions at specific concentrations of ligand and protein were determined from turbidity assays as described above (49). The ligand hub concentrations were chosen to cover as wide a range as possible. Cloud point temperature values were determined by fitting a Four Parameter Logistic Regression model to the data using MATLAB R2019b:

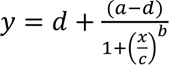

where *a* and *d* are minimum and maximum absorbance values, *b* is the Hill slope reflecting steepness of the phase transition and *c* is the temperature at the inflection point or the cloud point temperature (T_cp_).

T_cp_ values were used to define the coexistence curve as a function of ligand:protein ratio (ligand refers to Ub mutants/HT6-Ub/M1-Ub4 and protein to either UBQLN2 450C or full length UBQLN2). Here, results were averaged from data collected using a total of six trials with proteins from two separate protein preps.

### NMR Experiments

All NMR data were collected at 25°C using a Bruker Avance III 800 MHz spectrometer equipped with TCI cryoprobe. Protein solutions were prepared in a 20 mM NaPhosphate buffer (pH 6.8) with 0.5 mM EDTA, 0.02 % NaN_3_, and 5 % D_2_O. Data collected were processed using NMRPipe (50) and analyzed using CCPNMR 2.5.2 (51). Chemical shift perturbations (CSPs) were quantified as follows:

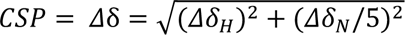

Here Δδ_*H*_ and Δδ_*N*_ are the differences in ^1^H and ^15^N chemical shifts in ppm, respectively.

### 15N Relaxation Experiments

Longitudinal (R_1_) and transverse (R_2_) ^15^N relaxation rates were measured for 200 µM samples of Ub and different HT6-Ub constructs using previously described protocols (48, 52). Relaxation inversion recovery periods for R_1_ experiments were 4 ms (*2), 600 ms (*2), and 1000 ms (*2), using an interscan delay of 2.5 s. Total spin-echo durations for R_2_ were 8 ms (*2), 32 ms (*2), 48 ms (*2), 64 ms (*2), and 80 ms (*2) using an interscan delay of 2.5 s. All relaxation experiments were acquired using spectral widths of 12 and 24 ppm in the ^1^H and ^15^N dimensions, respectively, with corresponding acquisition times of 110 ms and 31 ms. Relaxation rates were calculated by fitting peak heights to a mono-exponential decay using RELAXFIT (53). The average R_1_ and R_2_ values >63 residues in secondary structure elements of Ub were reported.

### NMR Titration Experiments and K_d_ Determination

^1^H-^15^N SOFAST-HMQC experiments were used for titrations of different Ub mutants/ HT6-Ub/ M1-Ub4 constructs into UBQLN2 UBA samples. Unlabeled protein ligand (Ub mutants/ HT6-Ub constructs/ M1-Ub4 constructs) was titrated into 50 µM samples of ^15^N-labeled protein (usually UBQLN2 UBA domain) and the binding was monitored as a function of different ligand:protein ratios. At each titration point, it was assumed that the CSP for each backbone amide was a weighted average between the free (Δδ=0) and ligand-bound states (Δδ= Δδ_*max*_). Therefore, the CSP reports on the relative population of the ligand-bound state, such that Δδ = Δδ_*max*_ ∗ [*PL*]/[*Pt*], where [PL] and [P_t_] represent the ligand-bound and the total UBA protein concentrations, respectively. Data fitting for each amide was performed using an in-house MATLAB program, with the assumption of a single-site binding model (1:1 stoichiometry). The equation for single-site binding for fast exchange is as follows:

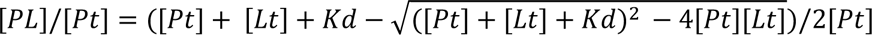

Here, [*Pt*] and [*Lt*] are total protein and ligand concentrations, respectively, and *K*_*d*_ is the binding affinity. Reported K_d_ values for each titration experiment were averages of residue-specific K_d_ values. The errors reflect the standard deviation of these values.

### SEC-MALS-SAXS Experiments

SAXS was performed at BioCAT (beamline 18ID at the Advanced Photon Source, Chicago) with in-line size exclusion chromatography (SEC-SAXS) to separate sample from aggregates and other contaminants thus ensuring optimal sample quality and multiangle light scattering (MALS), dynamic light scattering (DLS) and refractive index measurement (RI) for additional biophysical characterization (SEC-MALS-SAXS). Sample was loaded onto a Superdex 200 Increase 10/300 GL column (Cytiva) run by 1260 Infinity II HPLC (Agilent Technologies) at 0.6 ml/min. The flow passed through (in order) the Agilent UV detector, a MALS detector and a DLS detector (DAWN Helios II, Wyatt Technologies), and an RI detector (Optilab T-rEX, Wyatt). The flow then went through the SAXS flow cell. The flow cell consists of a 1.0 mm ID quartz capillary with ∼20 µm walls. A coflowing buffer sheath is used to separate the sample from the capillary walls, helping prevent radiation damage (54). Scattering intensity was recorded using a Pilatus3 X 1 M (dectris) detector which was placed 3.6 m from the sample giving access to a q-range of 0.003 Å^-1^ to 0.35 Å^-1^. 0.5 s exposure was acquired every 2 s during elution, and data were reduced using BioXTAS RAW 2.1.1 (55). Buffer blanks were created by averaging regions flanking the elution peak (see Fig S4, 7, 11) and subtracted from exposure selected from the elution peak to create the I(*q*) vs. *q* curves used for subsequent analysis. Molecular weights and hydrodynamic radii were calculated from the MALS and DLS data respectively using ASTRA 7 software (Wyatt). Additionally, R_g_ and I(0) values were obtained using the entire q-range of the data by calculating the distance distribution functions, P(*r*) vs. r, using GNOM (56). All SEC-MALS-SAXS parameters for data collection and analysis can be found in Table S5.

### Representative structure determination of ligand hubs from SAXS and SASSIE

We employed SASSIE (30) to generate structural ensembles for various HT6-Ub and M1-Ub4 ligand hubs with different linker lengths. Starting PDB structure files were built using AlphaFold2 (29). For M1-Ub4 ligands, we used the monomer configuration generator module of SASSIE to initially build 30,000 structures. As HT6-Ub ligands are tetrameric complexes, we used the Complex configuration generator module to build 30,000 structures. Monte Carlo moves about the ɸ/Ѱ backbone torsion angles were permitted only for selected residues (i.e., only these residues were deemed flexible in the structures) as denoted in Table S5, and each move was restricted to a maximum of 30 degrees. Trial structures were rejected if there were Cα-atom steric clashes within 3 Å. This yielded around 13,000 – 21,000 sterically-allowed structures (Table S6). X-ray scattering curves were then calculated for each of these structures. The single structure with the lowest χ^2^ (in comparison to the experimental X-ray scattering curve for the specific ligand hub) was then selected as the representative structure as shown in Fig. 2 and 4.

### Data availability

SAXS data for HT6-Ub & M1-Ub4 ligand hubs are deposited in SASBDB: https://www.sasbdb.org/project/1831/. All other data are available from corresponding authors upon request.

