## Supplementary Material for "Decoding optimal ligand design for multicomponent condensates"

*Table of Contents*

Theory for Ligand-mediated phase transitions
Figure S1
Figure S2
Figure S3
Figure S4
Figure S5
Figure S6
Figure S7
Figure S8
Figure S9
Figure S10
Figure S11

Table S1
Table S2
Table S3
Table S4
Table S5

### Theory for ligand-mediated phase transitions

#### Solution Free Energy

We have developed a theory to reconcile the observation that monoubiquitin and polyubiquitin inhibit and promote phase separation respectively. The first requirement of the model is that it must capture the binding equilibrium of ubiquitin and UBQLN2 into higher order complexes. We begin by writing down the free energy of the dilute state, which we denote with the index V = "vapor":

(Eq. S1)

$$F_V = c_{h,V}\left(\ln\frac{c_{h,V}}{c_0} - 1 + f_h\right) + c_{d,V}\left(\ln\frac{c_{d,V}}{c_0} - 1 + f_d\right) + \sum_{n=1}^{N} c_{n,V}\left(\ln\frac{c_{n,V}}{c_0} - 1 + f_n\right)$$
$$+ \mu_h(c_{H,V} - c_{h,V} - \sum_{n=1}^{N} c_{n,V}) + \mu_d(c_{D,V} - c_{d,V} - \sum_{n=1}^{N} n\, c_{n,V})$$

Here $c_{h,V}$ and $c_{d,V}$ are the concentrations of hub and driver molecules in the monomer state, respectively (i.e., not bound to any other hub or driver molecule). N is the number of driver molecules that a hub can bind (N=4 in the case of Ub4) and $c_{n,V}$ is the concentration of hubs that are bound to n driver molecules ("n-mers"). $f_i$ is the free energy of species i and $c_i\left(\ln\frac{c_i}{c_0} - 1\right)$ is the mixing entropy of that species, where $c_0$ is a reference concentration. The chemical potentials $\mu_h$ and $\mu_d$ serve as Lagrange multipliers to ensure that the concentration of molecules in the monomer and n-mer states add up to the total concentration of hubs, $c_{H,i}$, and driver molecules, $c_{D,i}$. (In our notation, the capital index (e.g. H, D) indicates the total concentration while the lowercase (e.g. h, d) indicates the monomer state).

The free energy of the dense state (denoted with the subscript L = "liquid") has a similar form:

(Eq. S2)

$$F_L = c_{h,L}\left(\ln\frac{c_{h,L}}{c_0} - 1 + f_h + s_h\right) + c_{d,L}\left(\ln\frac{c_{d,L}}{c_0} - 1 + f_d + s_d\right) + \sum_{n=1}^{N} c_{n,L}\left(\ln\frac{c_{n,L}}{c_0} - 1 + f_n + s_n\right)$$
$$+ \mu_h(c_{H,L} - c_{h,L} - \sum_{n=1}^{N} c_{n,L}) + \mu_d(c_{D,L} - c_{d,L} - \sum_{n=1}^{N} n\, c_{n,L})$$

Where $s_i$ is the energy to transfer species i from the dilute, vapor (V) state to the liquid (L) state (e.g. $s_h$ is the energy to transfer the hub from the vapor to liquid state).

#### Chemical and Partitioning Equilibrium

Minimizing $F_V$ and $F_L$ with respect to $c_i$ we find expressions for each of the species concentrations in the vapor phase:

(Eq. S3)

$$c_{h,V} = c_0 e^{-f_h+\mu_h}$$
$$c_{d,V} = c_0 e^{-f_d+\mu_d}$$
$$c_{n,V} = c_0 e^{-f_n+\mu_h+n\mu_d}$$

And in the liquid phase:

(Eq. S4)

$$c_{h,L} = c_0 e^{-f_h-s_h+\mu_h}$$
$$c_{d,L} = c_0 e^{-f_d-s_d+\mu_d}$$
$$c_{n,L} = c_0 e^{-f_n-s_n+\mu_h+n\mu_d}$$

The expressions for the monomers in the vapor phase can be rearranged to find expressions for the chemical potentials:

(Eq. S5)

$$\mu_h = \ln\frac{c_{h,V}}{c_0} + f_h$$
$$\mu_d = \ln\frac{c_{d,V}}{c_0} + f_d$$

These expressions can be used with Eqs. S3 and S4 to obtain the conditions for (unbound) monomer phase partitioning:

(Eq. S6)

$$c_{h,V} = c_{h,L} e^{s_h}$$
$$c_{d,V} = c_{d,L} e^{s_d}$$

Similarly, the chemical potentials can be substituted in the expressions for the n-mer concentrations:

(Eq. S7)

$$c_{n,V} = \frac{c_{h,V} c_{d,V}^n}{c_0^n} e^{-(f_n-f_h-n\,f_d)}$$
$$= \frac{c_{h,V} c_{d,V}^n}{k_{n,V}}$$

Where $k_{n,V}$ is the dissociation constant for the formation of n-mers in the dilute (vapor) phase.

Similarly, in the dense (liquid) phase we find:

(Eq. S8)

$$c_{n,L} = \frac{c_{h,L} c_{d,L}^n}{c_0^n} e^{-(f_n-f_h-n\,f_d)-(s_n-s_h-n\,s_d)}$$
$$= \frac{c_{h,L} c_{d,L}^n}{k_{n,L}}$$

where

(Eq. S9)

$$k_{n,L} = k_{n,V} e^{(s_n - s_h - n s_d)}$$
$$k_{n,L} = k_{n,V} e^{\Delta s}$$

Note that if the transfer free energy of an n-mer is given by the sum of its parts such that $s_n = s_h + n s_d$ then the dissociation constants are identical in each phase. Therefore, $\Delta s$, is a parameter that describes how oligomerization (hub-driver binding) modifies the interaction with the fluid (driver-only fluid). However, if the bonding constraints within the n-mer perturb the interactions with the surrounding phase, the oligomerization equilibrium will be different between the phases ($\Delta s$ will be non-zero).

The above formulation ensures that binding equilibria are satisfied in each phase and that the chemical potentials for each species are equal across phases.

Our next task is to determine a condition for the onset of phase separation. At the onset, we can consider the dense phase to be infinitesimally small so that all of the proteins are in the dilute (vapor) phase. Thus, $c_{H,V}$ and $c_{D,V}$ are equal to the total protein concentrations, which can be used with Eq. S7 to determine the monomer concentration ($c_{h,V}$ and $c_{d,V}$ (as described below)). These concentrations, in turn, can be used with Eq. S8 to find the monomer concentrations in the infinitesimal droplet. Next these concentrations are used with Eq. S9 to find the concentration of n-mers in the dense phase.

**Saturated Solution Condition**

To assess whether these concentrations represent a subsaturated or supersaturated solution we examine the total concentration of the driver molecules in the dense phase. When polymers phase separate, the mass concentration of the dense phase is insensitive to the molecular weight (or equivalently the polymerization number) of the individual molecules (57). This is because the microscopic interactions and mesh structure of the phase are both much smaller than the molecules. In agreement with this expectation, the concentration of UBQLN2 molecules in the dense phase is nearly constant regardless of whether the UBQLN2 are monomers or oligomerized by a hub (8). Therefore, we adopt, as the criteria for phase separation, the condition that the total concentration of UBQLN2 in the dense phase is equal to the concentration of the UBQLN2-only fluid (i.e. in a UBQLN2-only phase separating solution):

(Eq. S10)

$$c_{d,L} + \sum_n n\, c_{n,L} = c_{D,L}$$

where the total concentration $c_{D,L}$ is taken to be the constants $c_{D,L}$ = 10 mM for 450C and $c_{D,L}$ = 2 mM for full length UBQLN2 (8).

An important insight from Eq. S10 is that hub molecules facilitate phase separation by lowering the chemical potential of driver molecules in the dense phase. This is most easily seen by examining the unbound driver chemical potentials in the two limiting cases:

- In a sub-saturated solution, the UBQLN2 concentration will add up to less than the pure UBQLN2 fluid: $c_{d,L} + \sum_n n\, c_{n,L} < c_{D,L}$. In this case, the dense phase will collapse to fill the voids and optimize the UBQLN2-UBQLN2 contacts. After this collapse, the monomer (unbound driver) concentration in the dense phase will increase such that the chemical potential in the dense phase is greater than the dilute phase: $\ln\frac{c_{d,V}}{c_0} < \ln\frac{c_{d,L}}{c_0} + s_d$. This will drive the monomers (unbound drivers) to leave the dense phase, causing it to shrink.

- Conversely, in a supersaturated solution we would find $c_{d,L} + \sum_n n\, c_{n,L} > c_{D,L}$ implying that the driver molecules are packed too close together. In this case the dense phase will expand, lowering the concentration of monomers. Again comparing the resulting chemical potentials of the monomers we find $\ln\frac{c_{d,V}}{c_0} > \ln\frac{c_{d,L}}{c_0} + s_d$, which will drive additional monomers (unbound complexes) to enter the dense phase.

Therefore, the condition for the onset of phase separation is when Eq. S10 is satisfied and the monomer concentrations satisfy

(Eq. S11)

$$\ln\frac{c_{d,V}}{c_0} = \ln\frac{c_{d,L}}{c_0} + s_d$$

Which is equivalent to Eq. S6.

In the absence of hubs ($c_{D,L} = c_{d,L}$), this expression gives Eq. 3 from the main text:

(Eq. S12)

$$\frac{c_{sat}}{c_{D,L}} = e^{s_d}$$

**Temperature Dependence of Phase Separation**

The temperature dependence of $s_d$ is modeled by fitting the experimental cloud point temperatures to the quadratic function:

(Eq. S13)

$$c_{sat} = a(T - T_c)^2 + c_c$$

The best fit parameters are shown in Fig. S10. Combining this with our previous result (Eq. S12) we have:

$$e^{s_d} = \frac{c_{sat}}{c_{D,L}}$$

(Eq. S14)

$$e^{s_d} = \frac{a(T-T_c)^2 + c_c}{c_{D,L}}$$

Next, we express our condition for phase separation in terms of $s_d$:

(Eq. S15)

$$c_{D,L} = c_{d,L} + \sum_n n\, c_{n,L}$$

$$c_{D,L} = c_{d,L} + \sum_n n \frac{c_{h,L} c_{d,L}^n}{k_{n,L}}$$

Using Eqs. S6 and S9 this becomes

(Eq. S16)

$$c_{D,L} = c_{d,V} e^{-s_d} + c_{h,V} e^{-s_h} \sum_n n \frac{\left(c_{d,V} e^{-s_d}\right)^n}{k_{n,V} e^{\Delta s}}$$

$$1 = \frac{c_{d,V}}{c_{sat}} + \frac{c_{h,V} e^{-s_h}}{c_{D,L}} \sum_n n \frac{\left(\frac{c_{d,V} c_{D,L}}{c_{sat}}\right)^n}{k_{n,V} e^{\Delta s}}$$

Finally, to reduce the number of free parameters in the model, we make the approximation that $\Delta s$ is independent of n and obtain:

(Eq. S17)

$$1 = \frac{c_{d,V}}{c_{sat}} + \frac{c_{h,V} e^{-s_h - \Delta s}}{c_{D,L}} \sum_n n \frac{\left(\frac{c_{d,V} c_{D,L}}{c_{sat}}\right)^n}{k_{n,V}}$$

We refer to the quantity $s_h + \Delta s$ as the "inclusion energy". It accounts for two detrimental effects from transferring an n-mer to the dense phase. The first is the repulsive solvation energy of the hub and the second is the constraints preventing the bound drivers from attaining their optimal interactions with the fluid. This inclusion energy is the only free parameter in Eq. S17.

**Calculation of Cloud Point Curves**

Comparison between theory and experiment is complicated by the fact that Eq. S17 depends on the monomer (unbound) concentrations $c_d$ and $c_h$, which are not readily known from experiments. These quantities can be determined from the total concentrations as follows. First, we construct the partition function of a hub in the vapor phase:

(Eq. S18)

$$Q_h = 1 + \sum_{n=1}^{N} \frac{c_d^n}{k_{n,V}}$$

Where we approximate the equilibrium constants by assuming independent sites such that $k_{n,V} = k_0^n / {}_n^N C$, where $k_0$ is the dissociation constant between monoubiquitin and UBQLN2 at

25°C (Table S1) and ${}_{n}^{N}C$ is a binomial coefficient accounting for the multiplicity of binding. From the partition function we obtain an expression for the average number of bound drivers per hub

(Eq. S19)

$$\langle n \rangle = \frac{1}{Q_h} \sum_{n=1}^{N} n \frac{c_d^n}{k_{n,V}}$$

With this expression we can write an equation for $c_d$ in terms of the total concentrations:

(Eq. S20)

$$c_D = c_d + \langle n \rangle c_H$$

The left side of $c_d$ monotonically increases for positive values of $c_d$, meaning that there is only one root within the range $0 < c_d \leq c_D$ that can be readily found numerically. With $c_d$ known, $c_h$ is obtained from the statistical weight of the monomer term in the partition function

(Eq. S21)

$$c_h = \frac{c_H}{Q_h}$$

The quantities $c_h$ and $c_d$ are inserted into Eq. S17, along with Eq. S13, which is numerically solved for the onset temperature.

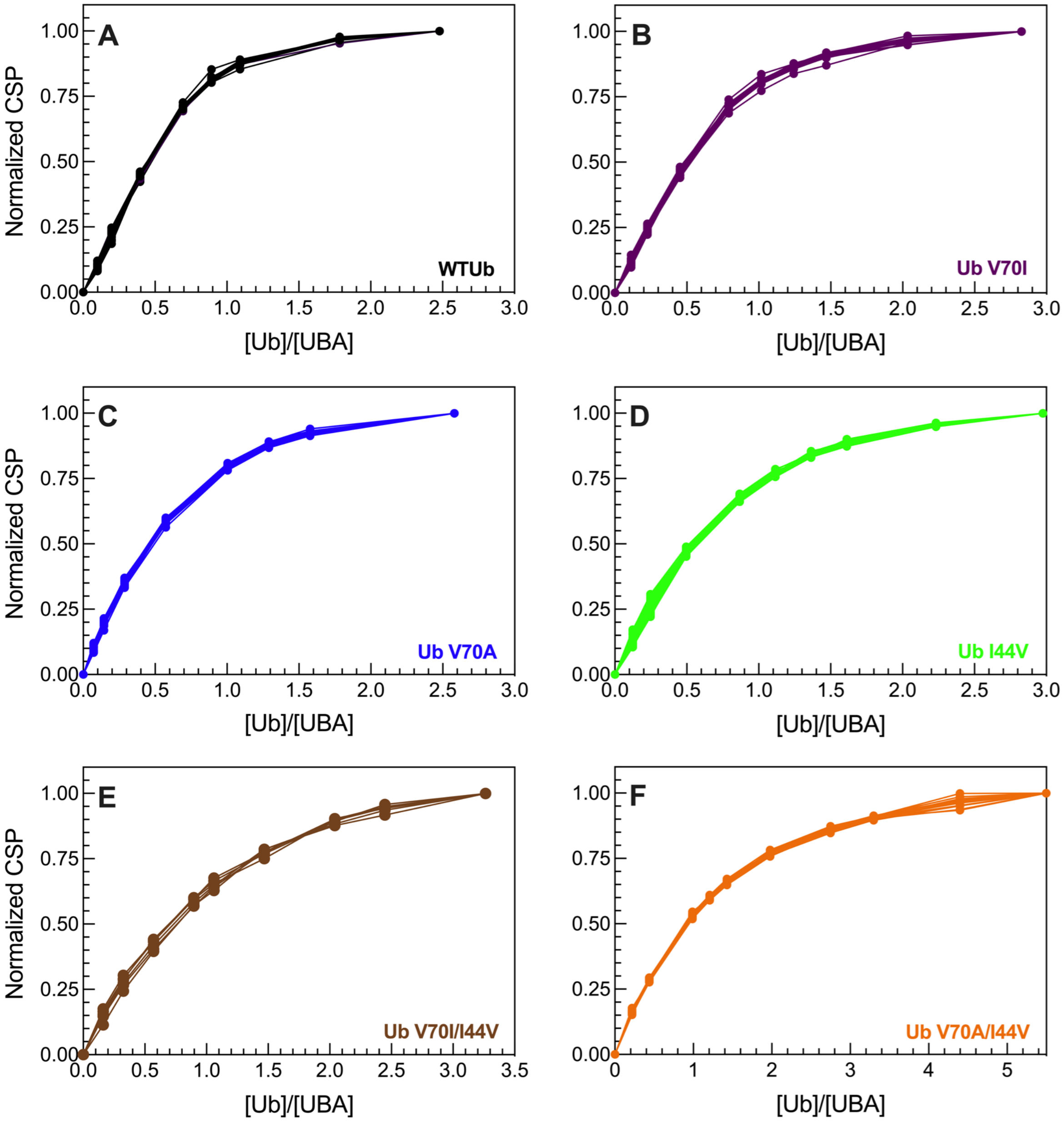


**Figure S1.** Residue-specific normalized backbone amide titration curves for 100 μM $^{15}$N UBQLN2 UBA resonances after titrating 0-250 μM of (A) WT Ub [22 amino acid resonances], (B) Ub V70I [21], (C) Ub V70A [17], (D) Ub I44V [23], (E) Ub V70I/I44V [6], & (F) Ub V70A/I44V [18]. See Table S1 for a list of specific amino acid resonances used.

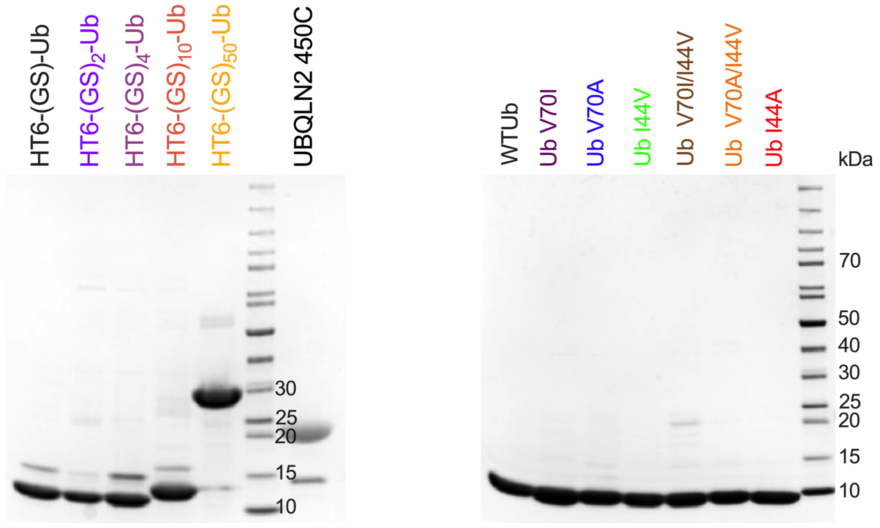


**Figure S2.** Representative Mini-PROTEAN TGX Precast gel images of (A) HT6-Ub and UBQLN2 450C, and (B) Ub mutants used in this study.

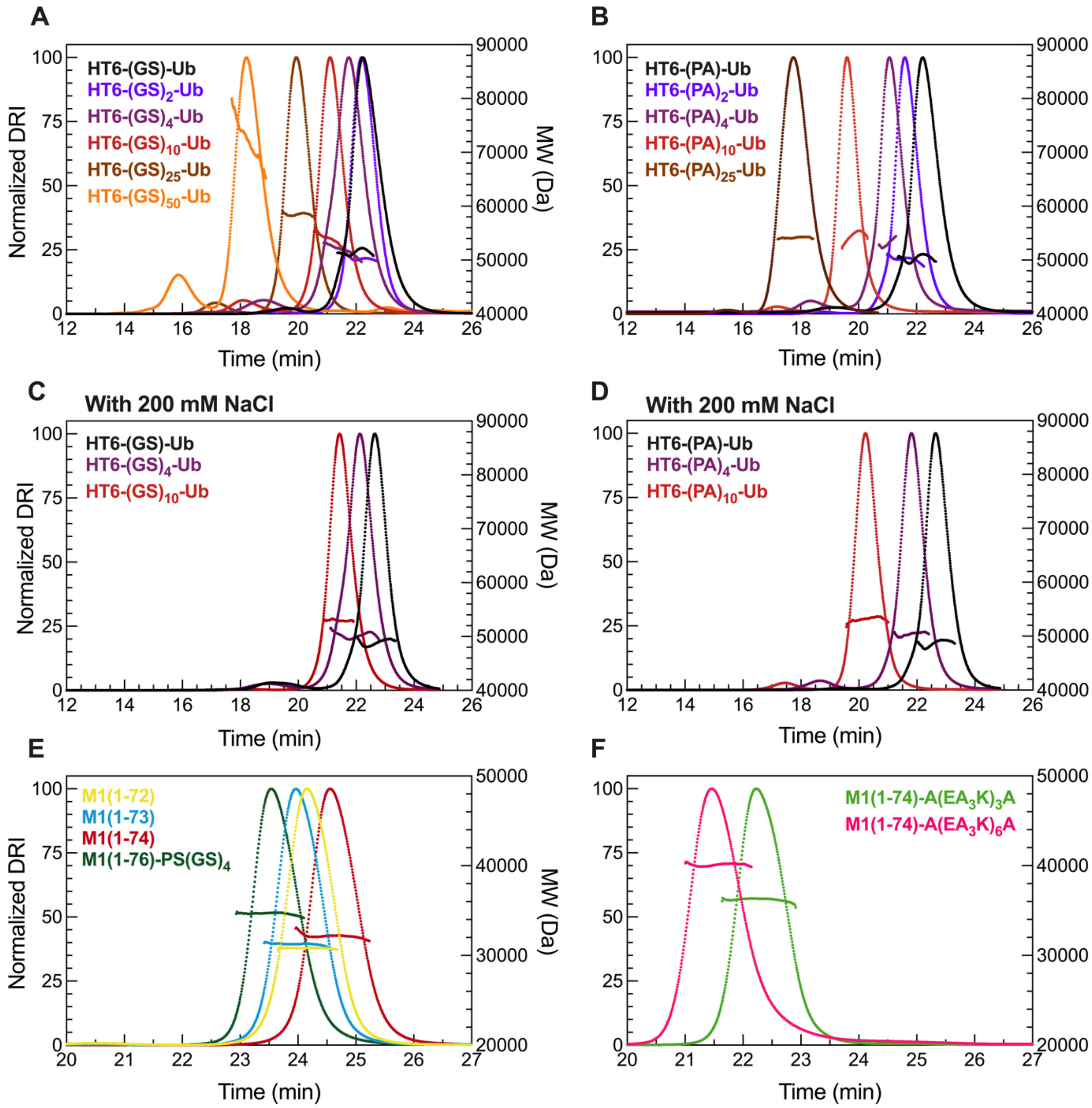


**Figure S3.** SEC-MALS profiles for various Ub4 ligand hubs used in this study. As expected, HT6-Ub is a tetramer. The molecular weight (MW) of (A) HT6-$(GS)_x$-Ub (B) HT6-$(PA)_x$-Ub (E) M1-linked Ub4 chains, and (F) M1-linked Ub4 chains with long Ub-Ub linkers were determined using SEC-MALS experiments. To test for the effect of salt on HT6-Ub oligomerization, we collected represented SEC-MALS profiles for (C) HT6-$(GS)_x$-Ub and (D) HT6-$(PA)_x$-Ub hubs. For all profiles collected above, the observed MW values are consistent with the expected MW of these constructs.

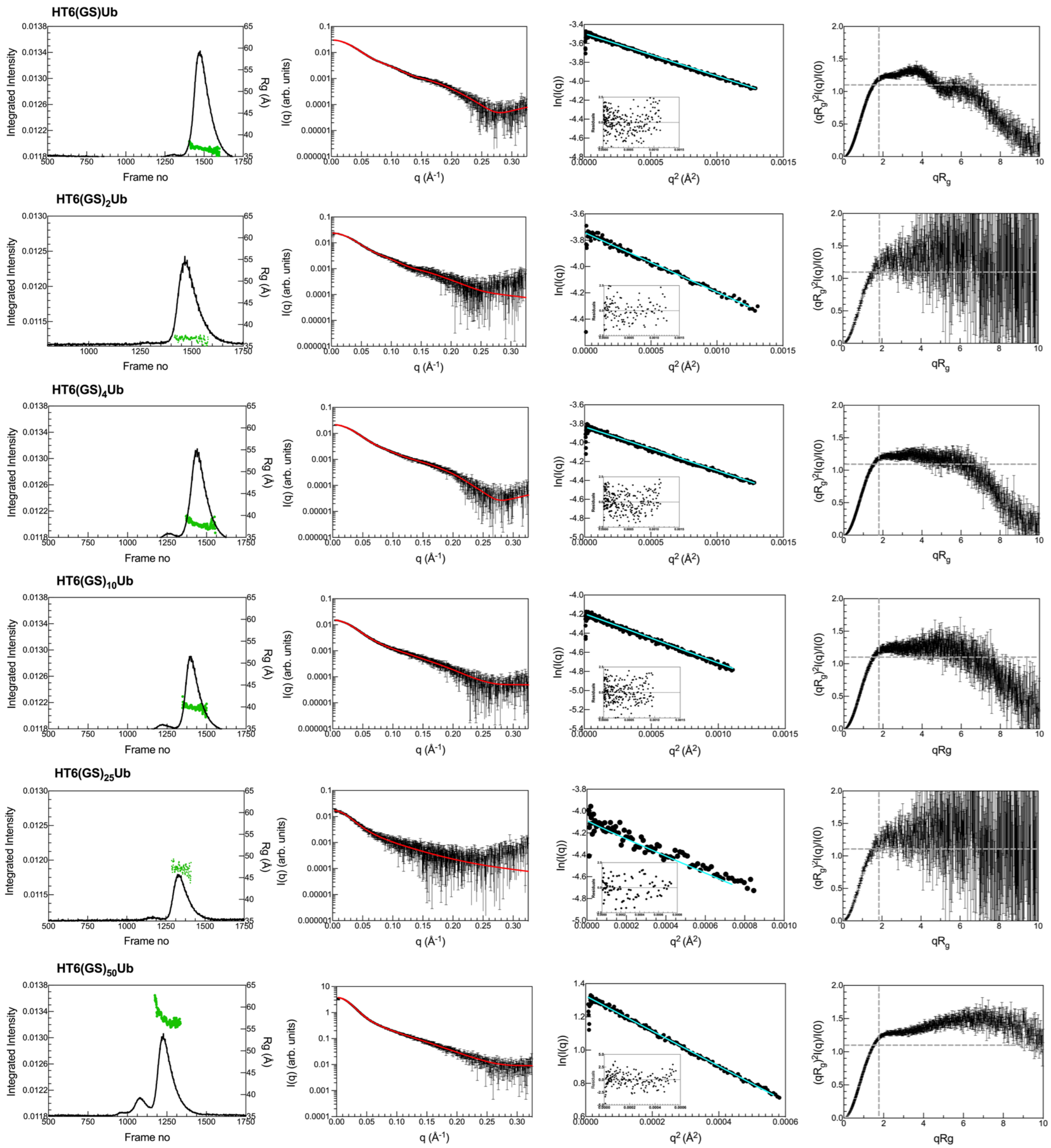


**Figure S4A.** (Left) SEC-SAXS profiles for HT6-(GS)-Ub, HT6-(GS)$_4$-Ub, HT6-(GS)$_{10}$-Ub, & HT6-(GS)$_{50}$-Ub. I(q) vs. q scattering curves (middle left) determined from frames (1438-1475) HT6-(GS)-Ub, (1453-1465) HT6-(GS)$_2$-Ub, (1448-1500) HT6-(GS)$_4$-Ub, (1423-1469) HT6-(GS)$_{10}$-Ub, (1322-1325) HT6-(GS)$_{25}$-Ub & (1165-1399) HT6-(GS)$_{50}$-Ub on the corresponding SEC-SAXS profiles (left). Red line in the I(q) profile indicates fit from P(r) analysis (see Fig. 2C). Cyan line in the Guinier plot (middle right) is the linear fit of ln(I(q)) vs. $q^2$, while inset shows residuals of fit. Dimensionless Kratky plots (right) include dashed lines to indicate where a globular protein would peak. Increase in polyUb chain flexibility from top (HT6-(GS)-Ub) to bottom (HT6-(GS)$_{50}$-

Ub) is indicated by shifts in the peak position up and to the right of the globular peak and larger plateaus in the higher q region.

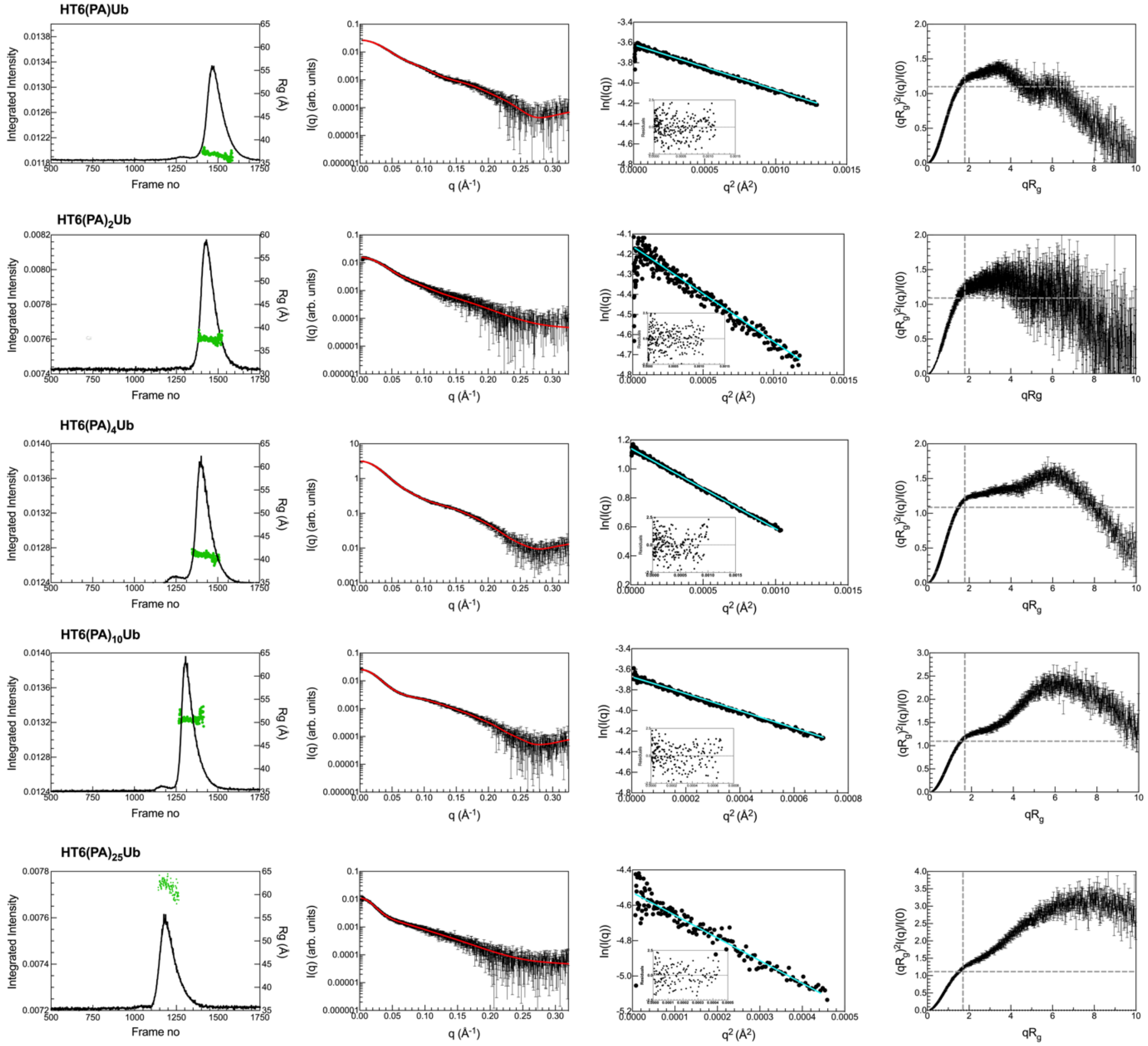


**Figure S4B.** SEC-SAXS profiles for HT6-(PA)-Ub, HT6-$(PA)_2$-Ub, HT6-$(PA)_4$-Ub, HT6-$(PA)_{10}$-Ub & HT6-$(PA)_{25}$-Ub. I(q) vs. q scattering curves (middle left) determined from frames (1478-1502) HT6-(PA)-Ub, (1425-1430) HT6-$(PA)_2$-Ub, (1452-1476) HT6-$(PA)_4$-Ub, (1352-1376) HT6-$(PA)_{10}$-Ub & (1183-1195) HT6-$(PA)_{25}$-Ub on the corresponding SEC-SAXS profiles (left). Red line in the I(q) profile indicates fit from P(r) analysis (see Fig. 2C). Cyan line in the Guinier plot (middle right) is linear fit of ln(I(q)) vs. $q^2$, while inset shows residuals of fit. Dimensionless Kratky plots (right) include dashed lines to indicate where a globular protein would peak. Increase in polyUb chain flexibility from top (HT6-(PA)-Ub) to bottom (HT6-$(PA)_{25}$-Ub) is indicated by shifts in the peak position up and to the right of the globular peak and larger plateaus in the higher q region.

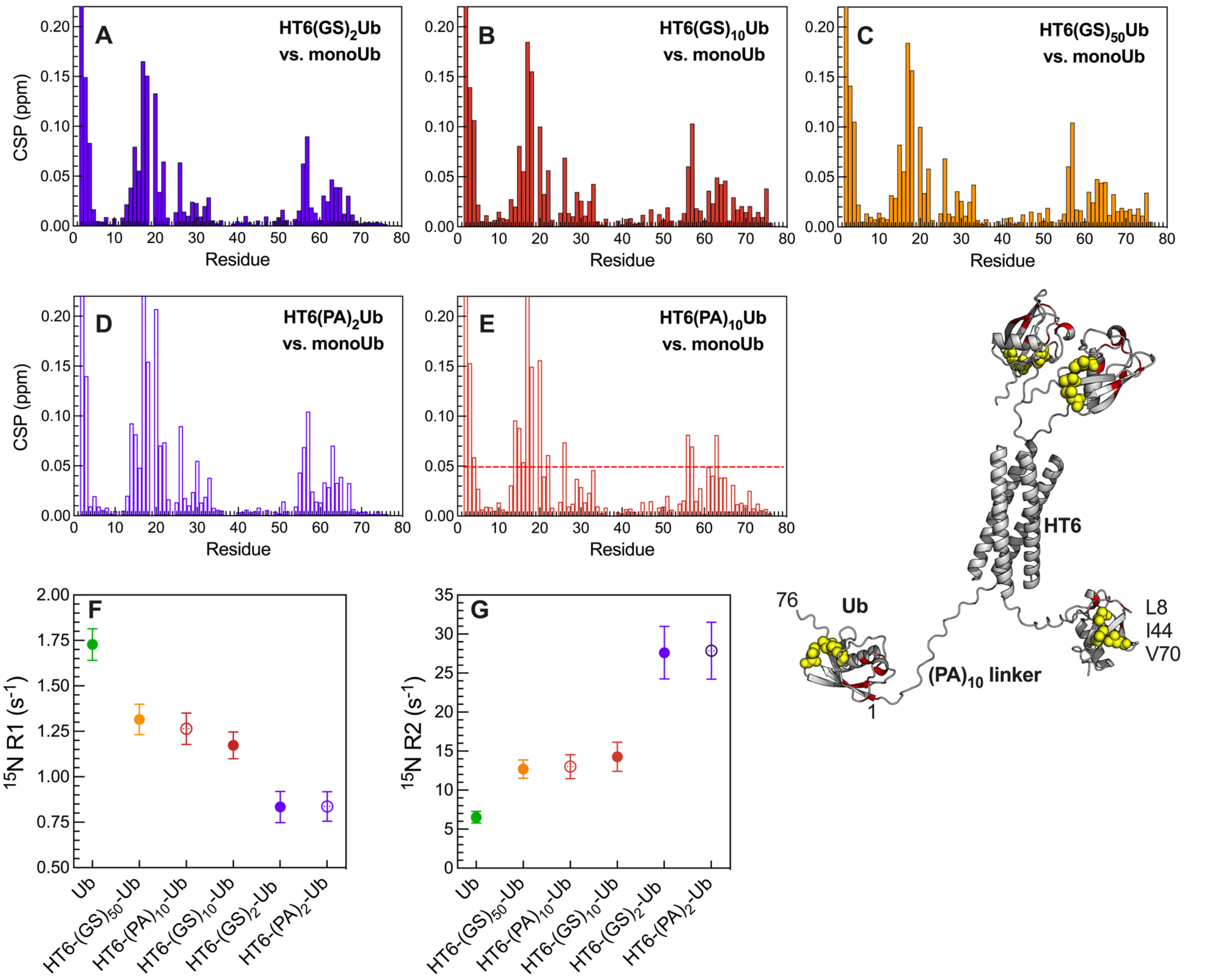


**Figure S5.** Comparison of Ub amide chemical shift perturbations (CSPs) for resonances in (A) HT6-(GS)$_2$-Ub, (B) HT6-(GS)$_{10}$-Ub, (C) HT6-(GS)$_{50}$-Ub, (D) HT6-(PA)$_2$-Ub, (E) HT6-(PA)$_{10}$-Ub vs. monoUb. CSPs > 0.05 ppm for (E) were mapped onto representative HT6-(PA)$_{10}$-Ub structure (see Fig. 2B) showing that only resonances spatially near the linker were perturbed, while not impacting the hydrophobic patch of Ub (yellow spheres). (F, G) Average $^{15}$N $R_1$ and $R_2$ relaxation rates for Ub resonances in secondary structure organized in terms of decreasing $R_1$ and increasing $R_2$ rates. Error bars denote standard deviation of residue-specific relaxation rates (>63 residues used) per construct.

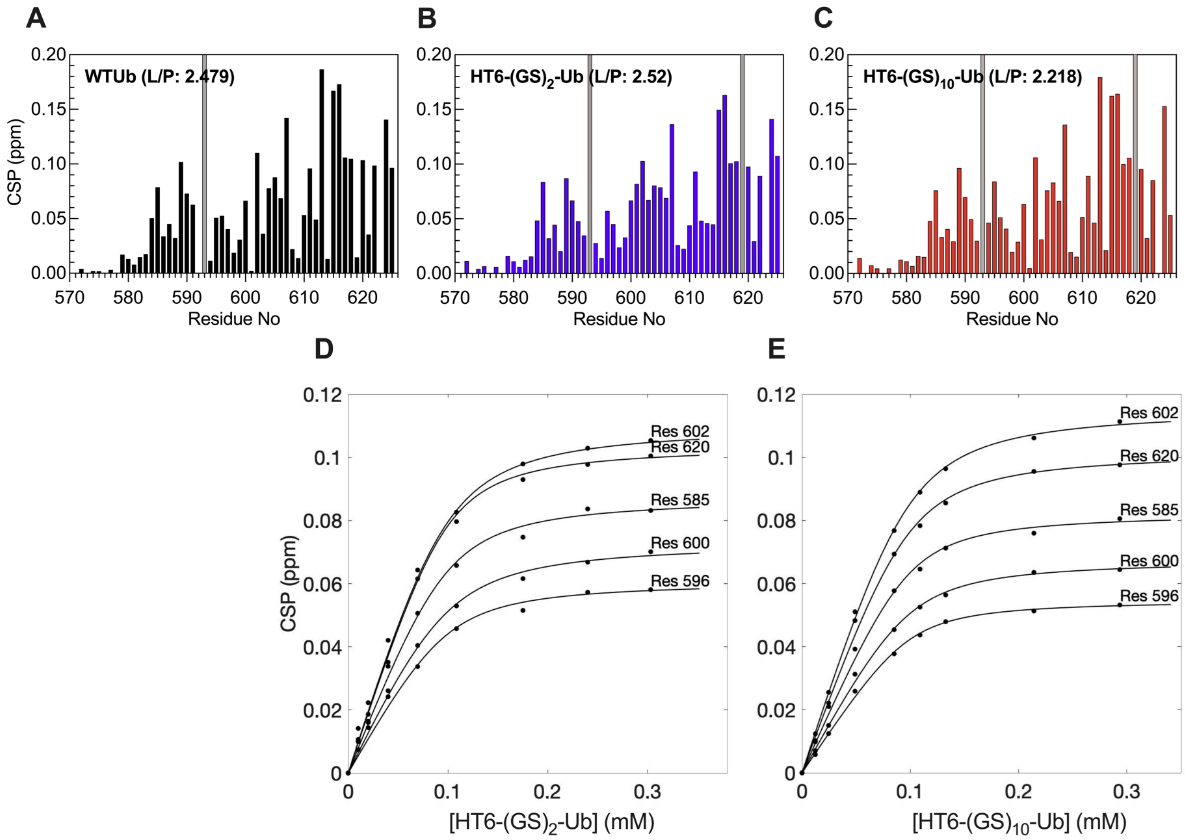


**Figure S6.** Comparison of backbone amide CSPs for UBA UBQLN2 (100 μM) upon titration with (A) WT Ub (ligand:protein (L:P=2.48), (B) HT6-$GS_2$-Ub (L:P=2.52) & (C) HT6-$GS_{10}$-Ub (L:P=2.22). Gray bars mark the resonances that are completely attenuated at the end of the titration. (D,E) Residue-specific amide titration curves of (D) HT6-$(GS)_2$-Ub, and (E) HT6-$(GS)_{10}$-Ub. Here fits from a single-site binding model were superpositioned on experimental data points.

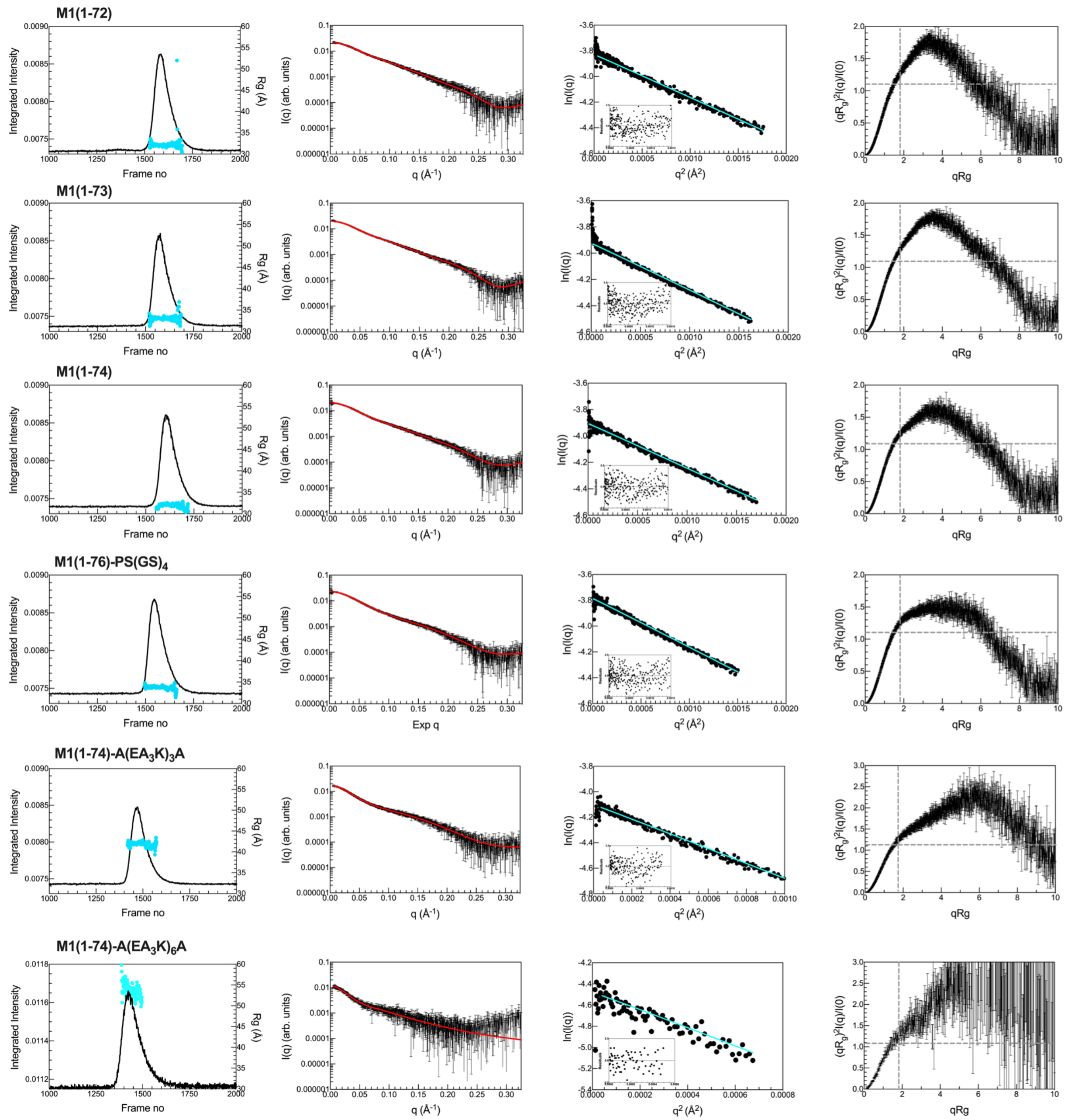


**Figure S7.** SEC-SAXS profiles for M1(1-72), M1(1-73), M1(1-74), M1(1-76)-PS(GS)$_4$, M1(1-74)-A(EA$_3$K)$_3$A & M1(1-74)-A(EA$_3$K)$_6$A. I(q) vs. q scattering curves (middle left) determined from frames (1580-1586) M1(1-72), (1556-1577) M1(1-73), (1592-1605) M1(1-74), (1540-1551) M1(1-76)-PS(GS)$_4$, (1440-1450) M1(1-74)-A(EA$_3$K)$_3$A, & (1437-1440) M1(1-74)-A(EA$_3$K)$_6$A on the corresponding SEC-SAXS profiles (left). Red line in the I(q) profile indicates fit from P(r) analysis (see Fig. 4). Cyan line in the Guinier plot (middle right) is linear fit of ln(I(q)) vs. $q^2$, while inset shows residuals of fit. Dimensionless Kratky plots (right) include dashed lines to indicate where a globular protein would peak.

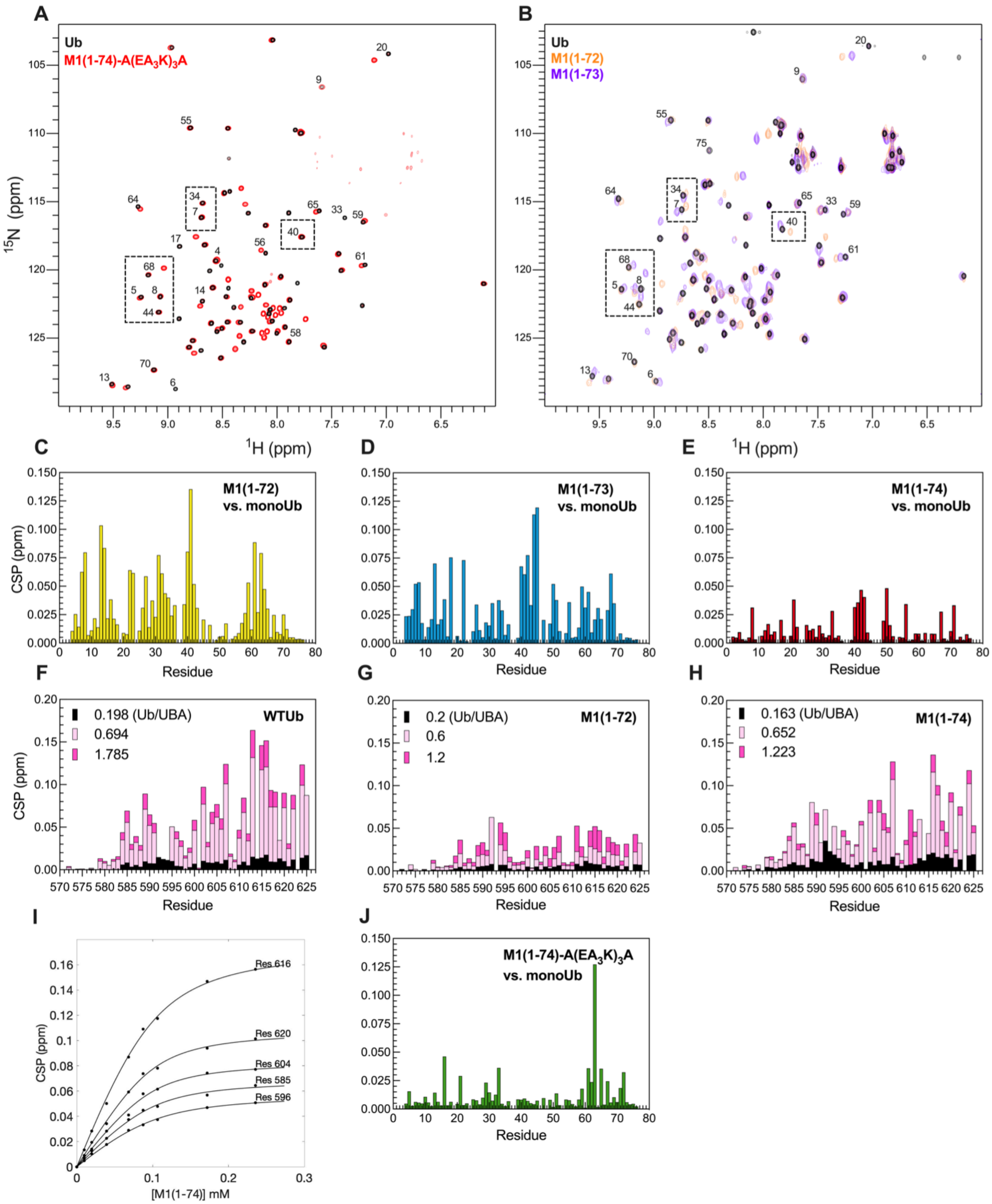


**Figure S8.** (A) $^{15}$N-$^{1}$H TROSY-HSQC spectra of 200 μM $^{15}$N labeled Ub (black) and M1-Ub4 (1-74)-A(EA$_3$K)$_3$A (red). (B) $^{15}$N-$^{1}$H SOFAST-HMQC spectra of 100 μM $^{15}$N labeled Ub (black), 200 μM M1-Ub4 (1-72) (orange), and 200 μM M1-Ub4 (1-73) (purple). Natural abundance spectra for M1-Ub4 (1-72) and M1-Ub4 (1-73) were collected with >1024 scans. Spectra were collected at 25 ˚C in 20 mM NaPhosphate buffer pH 6.8. (C, D, E) Chemical shift perturbations (CSPs) measured for backbone amide resonances in (C) M1-Ub4 (1-72), (D) M1-Ub4 (1-73), (E) M1-

Ub4 (1-74) with respect to $^{15}N$ labeled monoUb resonances under identical buffer conditions. (F, G, H) Amide CSPs measured at three different Ub:$^{15}N$ UBA ratios (as noted in figure) upon titrating unlabeled (F) WT Ub (G) M1-Ub4 (1-72) (H) M1-Ub4 (1-74) into 100 μM $^{15}N$ UBA UBQLN2. (I) Residue-specific backbone amide titration curves for UBA resonances as unlabeled M1-Ub4 (1-74) was titrated into 100 μM $^{15}N$ UBA UBQLN2. Single-site binding model (black line) was fit to residue-specific titration curves (data points) to obtain $K_d$ values. (J) Chemical shift perturbations (CSPs) measured for backbone amide resonances in M1(1-74)-A($EA_3K$)$_3$A with respect to $^{15}N$ labeled monoUb resonances under identical buffer conditions. As we observed multiple resonances for select Ub residues in M1(1-72), M1(1-73), M1(1-74) & M1(1-74)-A($EA_3K$)$_3$A, the largest CSP values for each residue are reported here.

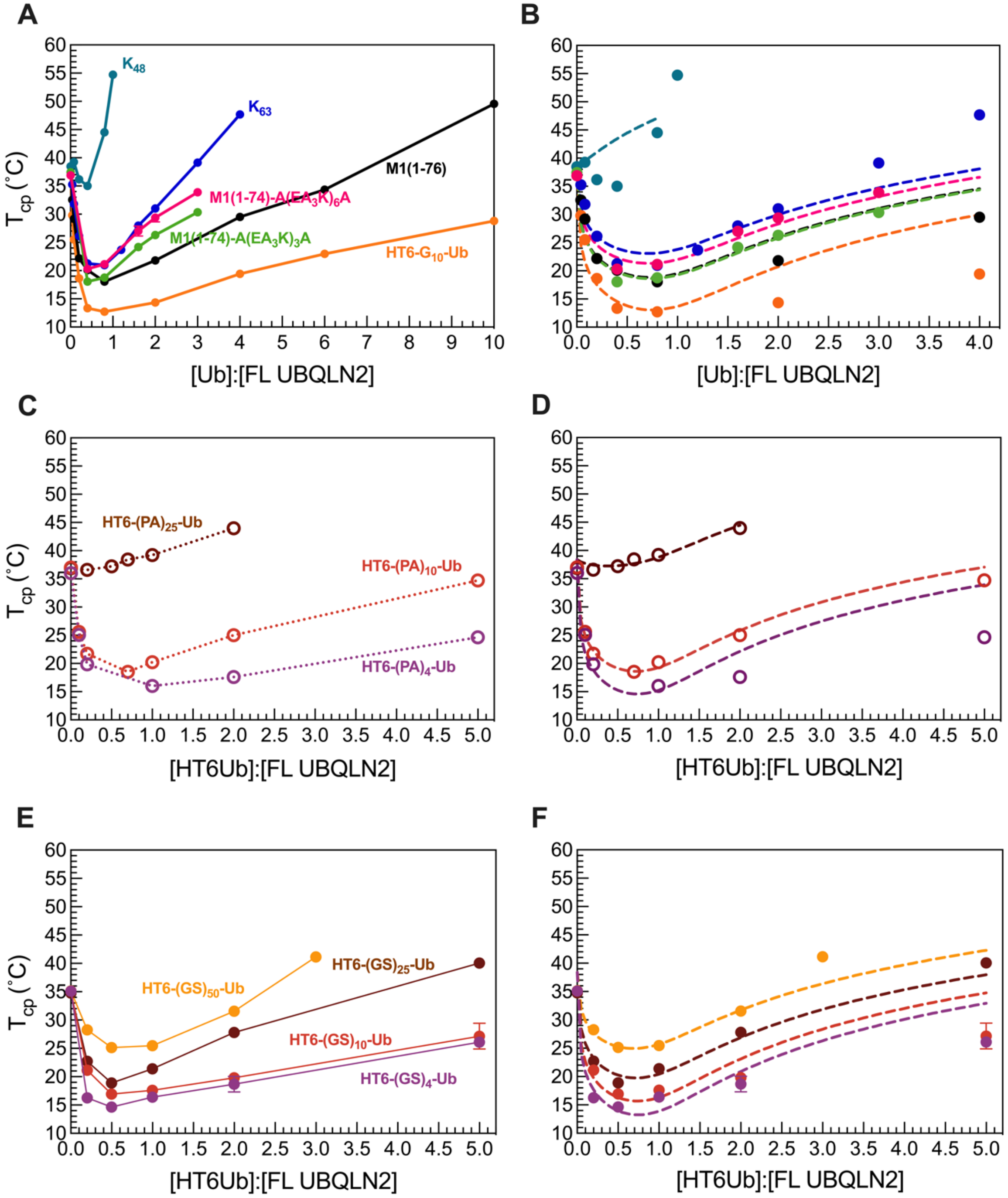


**Figure S9.** There is an optimal ligand architecture of polyUb that maximizes full-length (FL) UBQLN2 phase separation. (A, C) Experimental phase diagrams of FL UBQLN2 with natural and designed polyUb chains of different linkages. FL UBQLN2 concentration was kept constant at 30 µM. (B, D) Analytical theory fitted to experimental data.

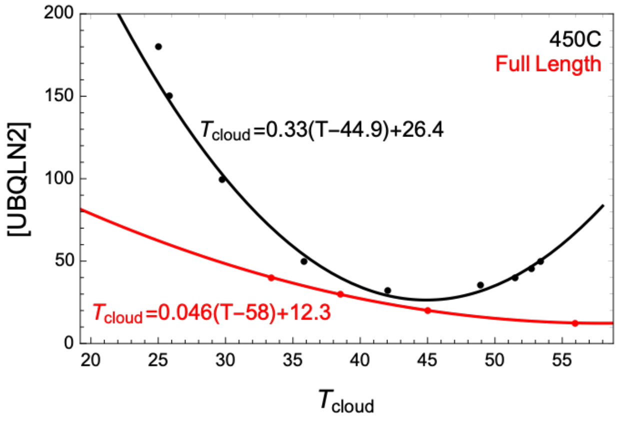


**Figure S10.** Comparison of experimental cloud point measurements (points) and fits to Eq. S13 for full length UBQLN2 (red) and 450C variant of UBQLN2 (black). Data for full length UBQLN2 and 450C variant of UBQLN2 are from (8) and (58), respectively.

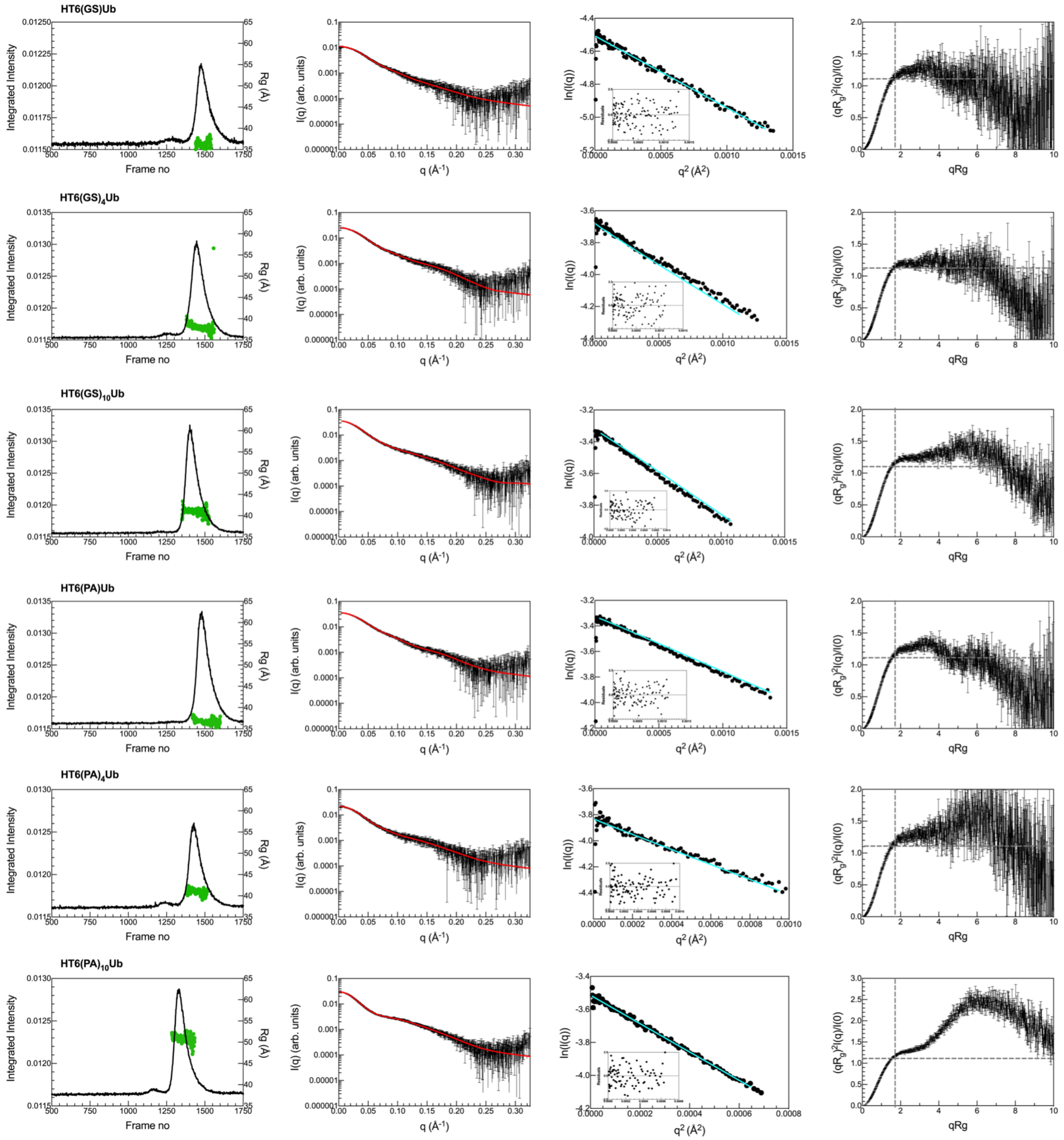


**Figure S11.** (A) SEC-SAXS profiles for HT6-(GS)-Ub, HT6-(GS)$_4$-Ub, HT6-(GS)$_{10}$-Ub, HT6-(PA)-Ub, HT6-(PA)$_4$-Ub, & HT6-(PA)$_{10}$-Ub collected in 20 mM phosphate buffer pH 6.8 with 200 mM NaCl, 0.5 mM EDTA & 0.02 % NaAz. I(q) vs. q scattering curves (middle left) determined from frames (1455-1513) HT6-(GS)-Ub, (1459-1482) HT6-(GS)$_4$-Ub, (1387-1408) HT6-(GS)$_{10}$-Ub, (1473-1484) HT6-(PA)-Ub, (1418-1428) HT6-(PA)$_4$-Ub & (1314-1354) HT6-(PA)$_{10}$-Ub on the corresponding SEC-SAXS profiles (left). Red line in the I(q) profile indicates fit from P(r) analysis. Cyan line in the Guinier plot (middle right) is the linear fit of ln(I(q)) vs. $q^2$, while inset shows residuals of fit. Dimensionless Kratky plots (right) include dashed lines to indicate where a globular protein would peak.

**Table S1.** Calculated average binding affinity ($K_d$) values of UBQLN2 UBA domain with WT Ub, Ub mutants, HT6-$(GS)_2$/$(GS)_{10}$-Ub and M1(1-74) used in the study. The errors represent the standard deviation of $K_d$ values determined from multiple NMR resonances as listed below (see Fig. S1).

| Protein | $K_d$ (μM) | Amino acid resonances used in $K_d$ determination |
|---|---|---|
| WT Ub | 2.6 ± 0.3 | 584, 585, 589, 590, 591, 596, 600, 602, 604, 605, 606, 607, 610, 611, 613, 615, 616, 617, 618, 620, 622, 624 |
| V70I | 6.0 ± 0.9 | 585, 589, 591, 593, 600, 602, 604, 605, 606, 607, 611, 614, 615, 616, 617, 618, 619, 620, 621, 622, 624 |
| V70A | 7.4 ± 0.6 | 589, 590, 596, 600, 602, 604, 605, 606, 607, 610, 612, 615, 616, 617, 618, 620, 624 |
| I44V | 13.6 ± 1.2 | 585, 589, 590, 593, 595, 596, 600, 602, 604, 605, 606, 607, 611, 612, 614, 615, 616, 617, 618, 619, 620, 622, 624 |
| V70I/I44V | 39 ± 3.6 | 588, 591, 596, 600, 606, 622 |
| V70A/I44V | 59 ± 1.8 | 589, 590, 596, 602, 604, 605, 606, 607, 610, 611, 612, 614, 615, 616, 618, 620, 622, 624 |
| I44A | > 1000 | |
| HT6-$(GS)_2$-Ub | 11.7 ± 2.1 | 585, 590, 596, 600, 602, 604, 605, 607, 610, 616, 617, 618, 619, 620, 622, 624 |
| HT6-$(GS)_{10}$-Ub | 9.0 ± 1.5 | 585, 589, 590, 591, 595, 596, 600, 602, 604, 605, 606, 607, 610, 611, 613, 615, 616, 617, 619, 620, 622, 624 |
| M1(1-74)-Ub4 | 15.6 ± 3.8 | 585, 596, 600, 602, 604, 606, 607, 615, 616, 617, 618, 620, 622, 624 |

**Table S2. Structural parameters of HT6-Ub and M1 series from SAXS data analysis.** Indicated in parentheses are the methods/software used for $R_g$ analysis. [a]$R_g$ and errors were determined from the linear fit of ln(I(q)) vs. $q^2$ and [b]$R_g$ and errors were determined from choosing multiple values of $D_{max}$. Data collected in 20 mM sodium phosphate buffer pH 6.8 with 0.5 mM EDTA and 0.02 % $NaN_3$.

| Construct | $R_g$ (Å)[a] (Guinier) [a] | $R_g$ (Å)[b] (GNOM) [b] | $D_{max}$ (Å) (GNOM) |
|---|---|---|---|
| HT6-(GS)-Ub | 37.10 ± 0.06 | 38.19 ± 0.09 | 140 |
| HT6-$(GS)_2$-Ub | 36.75 ± 0.17 | 38.08 ± 0.19 | 137 |
| HT6-$(GS)_4$-Ub | 37.65 ± 0.07 | 38.89 ± 0.10 | 143 |
| HT6-$(GS)_{10}$-Ub | 39.74 ± 0.12 | 41.37 ± 0.14 | 148 |
| HT6-$(GS)_{25}$-Ub | 47.02 ± 0.62 | 51.01 ± 0.66 | 193 |
| HT6-$(GS)_{50}$-Ub | 56.31 ± 0.09 | 58.70 ± 0.15 | 227 |
| HT6-(PA)-Ub | 36.98 ± 0.09 | 38.12 ± 0.08 | 129 |
| HT6-$(PA)_2$-Ub | 38.03 ± 0.24 | 39.88 ± 0.26 | 150 |
| HT6-$(PA)_4$-Ub | 40.80 ± 0.06 | 42.00 ± 0.06 | 144 |
| HT6-$(PA)_{10}$-Ub | 50.52 ± 0.13 | 52.29 ± 0.20 | 184 |
| HT6-$(PA)_{25}$-Ub | 62.61 ± 0.23 | 66.24 ± 0.36 | 249 |
| M1(1-72) | 32.03 ± 0.11 | 34.09 ± 0.15 | 125 |
| M1(1-73) | 33.09 ± 0.07 | 35.08 ± 0.12 | 128 |
| M1(1-74) | 32.21 ± 0.09 | 33.76 ± 0.11 | 122 |
| M1(1-76) | 32.42 ± 0.03 | 34.22 ± 0.21 | 140 |
| M1(1-76)-PS$(GS)_4$ | 34.04 ± 0.10 | 35.73 ± 0.17 | 139 |
| M1(1-74)-A$(EA_3K)_3$A | 42.06 ± 0.19 | 45.25 ± 0.40 | 182 |
| M1(1-74)-A$(EA_3K)_6$A | 51.84 ± 1.18 | 56.19 ± 1.42 | 212 |
| HT6-(GS)-Ub, 200 mM NaCl | 36.16 ± 0.18 | 37.75 ± 0.12 | 126 |
| HT6-$(GS)_4$-Ub, 200 mM NaCl | 37.77 ± 0.14 | 38.73 ± 0.14 | 128 |
| HT6-$(GS)_{10}$-Ub, 200 mM NaCl | 41.16 ± 0.13 | 42.32 ± 0.14 | 142 |
| HT6-(PA)-Ub, 200 mM NaCl | 36.85 ± 0.12 | 37.81 ± 0.14 | 124 |
| HT6-$(PA)_4$-Ub, 200 mM NaCl | 41.05 ± 0.26 | 42.67 ± 0.24 | 147 |
| HT6-$(PA)_{10}$-Ub, 200 mM NaCl | 50.78 ± 0.16 | 52.81 ± 0.21 | 190 |

**Table S3. Inclusion energy values for 450C UBQLN2 & FL UBQLN2 with polyUb ligands (related to Figure 4I, 4J).**

| Ligand | Inclusion energy (kT) | |
|---|---|---|
| | 450C UBQLN2 | FL UBQLN2 |
| HT6-(GS)-Ub | 9.43 | - |
| HT6-$(GS)_2$-Ub | 9.83 | - |
| HT6-$(GS)_4$-Ub | 10.07 | 6.67 |
| HT6-$(GS)_{10}$-Ub | 10.78 | 7.07 |
| HT6-$(GS)_{25}$-Ub | 11.47 | 7.79 |
| HT6-$(GS)_{50}$-Ub | 13.24 | 8.80 |
| HT6-(PA)-Ub | 9.65 | - |
| HT6-$(PA)_2$-Ub | 9.73 | - |
| HT6-$(PA)_4$-Ub | 10.16 | 6.89 |
| HT6-$(PA)_{10}$-Ub | 11.35 | 7.58 |
| HT6-$(PA)_{25}$-Ub | 14.13 | 11.84 |
| M1(1-73) | 10.89 | - |
| M1(1-74) | 9.78 | - |
| M1(1-76) | 10.01 | 7.62 |
| M1(1-76)-PS$(GS)_4$ | 9.81 | - |
| M1(1-74)-A$(EA_3K)_3$A | 10.52 | 7.58 |
| M1(1-74)-A$(EA_3K)_6$A | 11.15 | 8.08 |
| K48-Ub4 | - | 14.52 |
| K63-Ub4 | - | 8.42 |

**Table S4. Constructs used in the study**

| Construct | Sequence |
|---|---|
| HT6-$G_{10}$-Ub | MTLREIEELLRKIIEDSVRSVAELEDIEKWLKKIGGGGGGGGGG<br>MQIFVKTLTGKTITLEVEPSDTIENVKAKIQDKEGIPPDQQRLIFAGKQLEDGRTLSDYNIQ<br>KESTLHLVLRLRGG |
| HT6-$G_{10}$-Ub V70I | MTLREIEELLRKIIEDSVRSVAELEDIEKWLKKIGGGGGGGGGGMQIFVKTLTGKTITLEV<br>EPSDTIENVKAKIQDKEGIPPDQQRLIFAGKQLEDGRTLSDYNIQKESTLHLILRLRGG |
| HT6-$G_{10}$-Ub V70A | MTLREIEELLRKIIEDSVRSVAELEDIEKWLKKIGGGGGGGGGG<br>MQIFVKTLTGKTITLEVEPSDTIENVKAKIQDKEGIPPDQQRLIFAGKQLEDGRTLSDYNIQ<br>KESTLHLALRLRGG |
| HT6-$G_{10}$-Ub I44V | MTLREIEELLRKIIEDSVRSVAELEDIEKWLKKIGGGGGGGGGGMQIFVKTLTGKTITLEV<br>EPSDTIENVKAKIQDKEGIPPDQQRLVFAGKQLEDGRTLSDYNIQKESTLHLVLRLRGG |
| HT6-$G_{10}$-Ub I44A | MTLREIEELLRKIIEDSVRSVAELEDIEKWLKKIGGGGGGGGGGMQIFVKTLTGKTITLEV<br>EPSDTIENVKAKIQDKEGIPPDQQRLAFAGKQLEDGRTLSDYNIQKESTLHLVLRLRGG |
| HT6-$G_{10}$-Ub V70I/I44V | MTLREIEELLRKIIEDSVRSVAELEDIEKWLKKIGGGGGGGGGGMQIFVKTLTGKTITLEV<br>EPSDTIENVKAKIQDKEGIPPDQQRLVFAGKQLEDGRTLSDYNIQKESTLHLILRLRGG |
| HT6-$G_{10}$-Ub V70A/I44V | MTLREIEELLRKIIEDSVRSVAELEDIEKWLKKIGGGGGGGGGGMQIFVKTLTGKTITLEV<br>EPSDTIENVKAKIQDKEGIPPDQQRLVFAGKQLEDGRTLSDYNIQKESTLHLALRLRGG |
| HT6-GS-Ub | MTLREIEELLRKIIEDSVRSVAELEDIEKWLKKIGSMQIFVKTLTGKTITLEVEPSDTIENVK<br>AKIQDKEGIPPDQQRLIFAGKQLEDGRTLSDYNIQKESTLHLVLRLRGG |
| HT6-$(GS)_2$-Ub | MTLREIEELLRKIIEDSVRSVAELEDIEKWLKKIGSGSMQIFVKTLTGKTITLEVEPSDTIEN<br>VKAKIQDKEGIPPDQQRLIFAGKQLEDGRTLSDYNIQKESTLHLVLRLRGG |
| HT6-$(GS)_4$-Ub | MTLREIEELLRKIIEDSVRSVAELEDIEKWLKKIGSGSGSGSMQIFVKTLTGKTITLEVEPS<br>DTIENVKAKIQDKEGIPPDQQRLIFAGKQLEDGRTLSDYNIQKESTLHLVLRLRGG |
| HT6-$(GS)_{10}$-Ub | MTLREIEELLRKIIEDSVRSVAELEDIEKWLKKIGSGSGSGSGSGSGSGSGSGSGSMQIFVK<br>TLTGKTITLEVEPSDTIENVKAKIQDKEGIPPDQQRLIFAGKQLEDGRTLSDYNIQKESTLH<br>LVLRLRGG |
| HT6-$(GS)_{25}$-Ub | MTLREIEELLRKIIEDSVRSVAELEDIEKWLKKIGSGSGSGSGSGSGSGSGSGSGSGSG<br>SGSGSGSGSGSGSGSGSGSGSGSGSGSMQIFVKTLTGKTITLEVEPSDTIENVKAKIQDKE<br>GIPPDQQRLIFAGKQLEDGRTLSDYNIQKESTLHLVLRLRGG |
| HT6-$(GS)_{50}$-Ub | MTLREIEELLRKIIEDSVRSVAELEDIEKWLKKIGSGSGSGSGSGSGSGSGSGSGSGSG<br>SGSGSGSGSGSGSGSGSGSGSGSGSGSGSGSGSGSGSGSGSGSGSGSGSGSGSG<br>SGSGSGSGSGSGSGSGSGSGSGSMQIFVKTLTGKTITLEVEPSDTIENVKAKIQDKEGIPPD<br>QQRLIFAGKQLEDGRTLSDYNIQKESTLHLVLRLRGG |
| HT6-PA-Ub | MTLREIEELLRKIIEDSVRSVAELEDIEKWLKKIPAMQIFVKTLTGKTITLEVEPSDTIENVK<br>AKIQDKEGIPPDQQRLIFAGKQLEDGRTLSDYNIQKESTLHLVLRLRGG |
| HT6-$(PA)_2$-Ub | MTLREIEELLRKIIEDSVRSVAELEDIEKWLKKIPAPAMQIFVKTLTGKTITLEVEPSDTIEN<br>VKAKIQDKEGIPPDQQRLIFAGKQLEDGRTLSDYNIQKESTLHLVLRLRGG |

| | |
|---|---|
| HT6-(PA)$_4$-Ub | MTLREIEELLRKIIEDSVRSVAELEDIEKWLKKIPAPAPAPAPAMQIFVKTLTGKTITLEVEPSDTIENVKAKIQDKEGIPPDQQRLIFAGKQLEDGRTLSDYNIQKESTLHLVLRLRGG |
| HT6-(PA)$_{10}$-Ub | MTLREIEELLRKIIEDSVRSVAELEDIEKWLKKIPAPAPAPAPAPAPAPAPAPAPAPAMQIFVKTLTGKTITLEVEPSDTIENVKAKIQDKEGIPPDQQRLIFAGKQLEDGRTLSDYNIQKESTLHLVLRLRGG |
| HT6-(PA)$_{25}$-Ub | MTLREIEELLRKIIEDSVRSVAELEDIEKWLKKIPAPAPAPAPAPAPAPAPAPAPAPAPAPAPAPAPAPAPAPAPAPAPAPAPAPAPAPAPAMQIFVKTLTGKTITLEVEPSDTIENVKAKIQDKEGIPPDQQRLIFAGKQLEDGRTLSDYNIQKESTLHLVLRLRGG |
| M1-Ub4 (1-72) | MQIFVKTLTGKTITLEVEPSDTIENVKAKIQDKEGIPPDQQRLIFAGKQLEDGRTLSDYNIQKESTLHLVLRMQIFVKTLTGKTITLEVEPSDTIENVKAKIQDKEGIPPDQQRLIFAGKQLEDGRTLSDYNIQKESTLHLVLRMQIFVKTLTGKTITLEVEPSDTIENVKAKIQDKEGIPPDQQRLIFAGKQLEDGRTLSDYNIQKESTLHLVLRMQIFVKTLTGKTITLEVEPSDTIENVKAKIQDKEGIPPDQQRLIFAGKQLEDGRTLSDYNIQKESTLHLVLRLRGG |
| M1-Ub4 (1-73) | MQIFVKTLTGKTITLEVEPSDTIENVKAKIQDKEGIPPDQQRLIFAGKQLEDGRTLSDYNIQKESTLHLVLRLMQIFVKTLTGKTITLEVEPSDTIENVKAKIQDKEGIPPDQQRLIFAGKQLEDGRTLSDYNIQKESTLHLVLRLMQIFVKTLTGKTITLEVEPSDTIENVKAKIQDKEGIPPDQQRLIFAGKQLEDGRTLSDYNIQKESTLHLVLRLMQIFVKTLTGKTITLEVEPSDTIENVKAKIQDKEGIPPDQQRLIFAGKQLEDGRTLSDYNIQKESTLHLVLRLRGG |
| M1-Ub4 (1-74) | MQIFVKTLTGKTITLEVEPSDTIENVKAKIQDKEGIPPDQQRLIFAGKQLEDGRTLSDYNIQKESTLHLVLRLRMQIFVKTLTGKTITLEVEPSDTIENVKAKIQDKEGIPPDQQRLIFAGKQLEDGRTLSDYNIQKESTLHLVLRLRMQIFVKTLTGKTITLEVEPSDTIENVKAKIQDKEGIPPDQQRLIFAGKQLEDGRTLSDYNIQKESTLHLVLRLRMQIFVKTLTGKTITLEVEPSDTIENVKAKIQDKEGIPPDQQRLIFAGKQLEDGRTLSDYNIQKESTLHLVLRLRGG |
| M1-Ub4 (1-76) | MQIFVKTLTGKTITLEVEPSDTIENVKAKIQDKEGIPPDQQRLIFAGKQLEDGRTLSDYNIQKESTLHLVLRLRGGMQIFVKTLTGKTITLEVEPSDTIENVKAKIQDKEGIPPDQQRLIFAGKQLEDGRTLSDYNIQKESTLHLVLRLRGGMQIFVKTLTGKTITLEVEPSDTIENVKAKIQDKEGIPPDQQRLIFAGKQLEDGRTLSDYNIQKESTLHLVLRLRGGMQIFVKTLTGKTITLEVEPSDTIENVKAKIQDKEGIPPDQQRLIFAGKQLEDGRTLSDYNIQKESTLHLVLRLRGG |
| M1(1-76)-PS(GS)$_4$<br><br>(linker in orange) | MQIFVKTLTGKTITLEVEPSDTIENVKAKIQDKEGIPPDQQRLIFAGKQLEDGRTLSDYNIQKESTLHLVLRLRGGPSGSGSGSGSMQIFVKTLTGKTITLEVEPSDTIENVKAKIQDKEGIPPDQQRLIFAGKQLEDGRTLSDYNIQKESTLHLVLRLRGGPSGSGSGSGSMQIFVKTLTGKTITLEVEPSDTIENVKAKIQDKEGIPPDQQRLIFAGKQLEDGRTLSDYNIQKESTLHLVLRLRGGPSGSGSGSGSMQIFVKTLTGKTITLEVEPSDTIENVKAKIQDKEGIPPDQQRLIFAGKQLEDGRTLSDYNIQKESTLHLVLRLRGG |
| M1(1-74)-A(EA$_3$K)$_3$A<br><br>(linker in orange) | MQIFVKTLTGKTITLEVEPSDTIENVKAKIQDKEGIPPDQQRLIFAGKQLEDGRTLSDYNIQKESTLHLVLRLRAEAAAKEAAAKEAAAKAMQIFVKTLTGKTITLEVEPSDTIENVKAKIQDKEGIPPDQQRLIFAGKQLEDGRTLSDYNIQKESTLHLVLRLRAEAAAKEAAAKEAAAKAMQIFVKTLTGKTITLEVEPSDTIENVKAKIQDKEGIPPDQQRLIFAGKQLEDGRTLSDYNIQKESTLHLVLRLRAEAAAKEAAAKEAAAKAMQIFVKTLTGKTITLEVEPSDTIENVKAKIQDKEGIPPDQQRLIFAGKQLEDGRTLSDYNIQKESTLHLVLRLRGG |
| M1(1-74)-A(EA$_3$K)$_6$A<br><br>(linker in orange) | MQIFVKTLTGKTITLEVEPSDTIENVKAKIQDKEGIPPDQQRLIFAGKQLEDGRTLSDYNIQKESTLHLVLRLRAEAAAKEAAAKEAAAKEAAAKEAAAKEAAAKAMQIFVKTLTGKTITLEVEPSDTIENVKAKIQDKEGIPPDQQRLIFAGKQLEDGRTLSDYNIQKESTLHLVLRLRAEAAAKEAAAKEAAAKEAAAKEAAAKEAAAKAMQIFVKTLTGKTITLEVEPSDTIENVKAKIQDKEGIPPDQQRLIFAGKQLEDGRTLSDYNIQKESTLHLVLRLRAEAAAKEAAAKEAAAKEAAAKEAAAKEAAAKA |

| | |
|---|---|
| | MQIFVKTLTGKTITLEVEPSDTIENVKAKIQDKEGIPPDQQRLIFAGKQLEDGRTLSDYNIQKE STLHLVLRLRGG |
| UBQLN2 450C | MRAMQALMQIQQGLQTLATEAPGLIPSFTPGVGVGVLGTAIGPVGPVTPIGPIGPIVPFTP IGPIGPIGPTGPAAPPGSTGSGGPTGPTVSSAAPSETTSPTSESGPNQQFIQQMVQALA GANAPQLPNPEVRFQQQLEQLNAMGFLNREANLQALIATGGDINAAIERLLGSQPSW |

**Table S5. SEC-MALS-SAXS Data Collection and Analysis for all Ub hubs in this study**

- See attached Excel file.

**Table S6. SASSIE parameters and results for generation of ligand hub conformational ensembles**

| Ligand hub | Flexible component of starting structure | Structures Generated | Accepted | Best single structure (reduced $\chi^2$) | Number of structures with lowest $\chi^2$ |
|---|---|---|---|---|---|
| HT6-$(PA)_4$-Ub | 35-42, 115-118 | 30000 | 20613 | 2.77 | 83<br>$\chi^2 < 4$ |
| HT6-$(PA)_{10}$-Ub | 35-54,126-129 | 30000 | 18786 | 1.06 | 13<br>$\chi^2 < 1.5$ |
| M1-Ub4 (1-72) | 72-73, 144-145, 216-217, 288-292 | 30000 | 13449 | 0.86 | 44<br>$\chi^2 < 1.0$ |
| M1-Ub4 (1-74) | 72-74, 146-148, 220-222, 294-298 | 30000 | 18927 | 1.03 | 47<br>$\chi^2 < 1.5$ |
| M1-Ub4 (1-74) $A(EA_3K)_6A$ | 72-76, 105-107, 178-182, 211-213, 284-288, 317-319, 390-394 | 30000 | 19359 | 0.63 | 140<br>$\chi^2 < 0.75$ |
